# Phylogenetic distribution and expression pattern analyses identified a divergent basal body assembly protein involved in land plant spermatogenesis

**DOI:** 10.1101/2021.07.25.453666

**Authors:** Shizuka Koshimizu, Naoki Minamino, Tomoaki Nishiyama, Emiko Yoro, Kazuo Ebine, Keiko Sakakibara, Takashi Ueda, Kentaro Yano

## Abstract

Oogamy is a form of sexual reproduction and evolved independently in animals, fungi, and plants. In streptophyte plants, Charophyceae, Coleochaetophyceae, bryophytes, lycophytes, ferns (monilophytes), and some gymnosperms (Cycads and Ginkgo) utilize spermatozoids as the male gamete. Plant spermatozoids commonly possess characteristic structures such as the spline, which consists of a microtubule array, the multilayered structure (MLS) in which the uppermost layer is continuum of the spline, and multiple flagella. However, the molecular mechanisms underpinning plant spermatogenesis remain to be elucidated. To identify the genes involved in plant spermatogenesis, we performed computational analyses and successfully found deeply divergent *BLD10*s by combining multiple methods and omics-data. We then validated the functions of candidate genes in the liverwort *Marchantia polymorpha* and the moss *Physcomitrium patens* and found that Mp*BLD10* and Pp*BLD10* are required for normal basal body and flagella formation. Mp*bld10* mutants exhibited defects in remodeling of the cytoplasm and nucleus during spermatozoid formation, thus Mp*BLD10* should be involved in chromatin reorganization and elimination of the cytoplasm during spermiogenesis. Streptophyte BLD10s are orthologous to BLD10/CEP135 family proteins, which function in basal body assembly, but we found that BLD10s evolved especially fast in land plants and MpBLD10 might obtain additional functions in spermatozoid formation through the fast molecular evolution. This study provides a successful example of combinatorial study from evolutionary and molecular genetic perspectives that elucidated a function of the key protein of the basal body formation that fast evolved in land plants.

## Introduction

Oogamy is a form of sexual reproduction using female and male gametes. The female gamete (egg cell) is non-motile and larger than the male gamete, whereas male gametes (sperm) are motile and smaller than female gametes. Oogamy evolved independently in animals, fungi, and plants (Spratt 1971; Simpson 2018), and it is a big question what genes drove evolution to oogamy, i.e., sperm production.

In streptophyte plants, sexual reproduction in Charophyceae, Coleochaetophyceae, and land plants are via oogamy. Among these organisms, Charophyceae, Coleochaetophyceae, bryophytes, lycophytes, ferns (monilophytes), and some gymnosperms (cycads and ginkgo) utilize spermatozoids as the male gamete. Oogamy in streptophyte plants is presumed to have originated from a single ancestor, then flagella of spermatozoids were lost independently in angiosperms, gymnosperms (in the common ancestor of cupressophytes, gnetophytes and Pinaceae), and Zygnematophyceae (Hodges et al. 2012). Plant spermatozoids commonly possess characteristic structures such as the spline, which consists of a microtubule array, the multilayered structure (MLS) in which the uppermost layer is a continuum of the spline and basal bodies are located on it, and multiple flagella. For decades, these structural features of spermatozoids have been investigated mainly by transmission electron microscopy (Norstog 1967; Carothers and Kreitner 1968; Kreitner and Carothers 1976; Graham and McBride 1979; Carothers and Duckett 1980; Renzaglia et al. 1985; Renzaglia and Duckett 1987). As reviewed by Renzaglia and Garbary (2001), in the spermatozoids of Charophyceae, bryophytes, and ferns, after the MLS develops, the nucleus becomes compacted and helically elongated along the spline, during which a major part of the cytoplasm is eliminated. Unlike in these species, in gymnosperms, the nucleus is neither condensed nor elongated, and the cytoplasm is not eliminated. The number of flagella in a spermatozoid of bryophytes or lycophytes is two, but a spermatozoid of ferns forms 20 - 50 or more flagella. Gymnosperms spermatozoids possess 1,000 - 50,000 flagella. Although morphological studies have been well conducted, the molecular and genetic players in plant spermatogenesis remain to be identified.

Currently, in addition to angiosperms, the genome sequences of a variety of streptophytes have been determined by progress of sequencing technologies (Rensing et al. 2008; Banks et al. 2011; Bowman et al. 2017; Li et al. 2018; Nishiyama et al. 2018; Zhao et al. 2019; Li et al. 2020; Wang et al. 2020), and a vast amount of omics data such as transcriptome have been accumulating in an online database, the Sequence Read Archive (SRA; Leinonen et al. 2010). Banks et al. (2011) reported that, after gene clustering, 32 of 137 ‘angiospermLoss’ groups (defined as present in at least two of the following: Chlamydomonas [*Chlamydomonas reinhardtii*], Physcomitrella [*Physcomitrium patens*], and Selaginella [*Selaginella moellendorffii*] but not in 15 angiosperms) harbored genes exhibiting similarity to flagella or basal body-related genes, consistent with the presence of flagellated cells in the three organisms. We envisioned that combining the phylogenetic distribution and expression data would yield a more specific set that could test their function using molecular genetic methods. We selected candidate genes specifically expressed in male reproductive tissues of Marchantia (*Marchantia polymorpha*) and Physcomitrella but excluded genes apparently present in Chlamydomonas to obtain a set that is worth investigating for the function in plant spermatogenesis. The functions of the candidate genes were examined in the liverwort *Marchantia polymorpha* and the moss *Physcomitrium* [*Physcomitrella*] *patens* (Rensing et al. 2020), for which molecular genetic techniques have been established (Schaefer 1997; Nishiyama et al. 2000; Ishizaki et al. 2008; Kubota et al. 2013). Loss-of-function mutants of the candidates Mp*BLD10* and Pp*BLD10* exhibited defects in basal body and flagella formation during spermatogenesis, suggesting that these genes are required for normal basal body and flagella formation. *BLD10*s were found to be putative orthologs of *BLD10* in *Chlamydomonas reinhardtii* (Cr*BLD10*), the product is required for assembly of the basal body. However, *BLD10*s evolved fast in the land plant lineage, and loss-of-function mutations in Mp*BLD10* and Pp*BLD10* resulted in different phenotypes in the liverwort and moss, which were also distinct from the phenotype in the chlorophyte Cr*bld10* mutant. Defects in reorganizing the cytoplasm and nucleus during spermatozoid formation in Mpbld10 mutants suggested that MpBLD10 plays a role in spermatozoid formation, in addition to the basal body formation. Thus, we present the results of a successful combinatorial study encompassing phylogenetic distribution, gene expression, and molecular genetics approaches, which unraveled that the function of the key component of the basal body formation is diverged during plant evolution.

## Results

### Four protein family groups were selected as candidates involved in spermatogenesis

To identify genes involved in plant spermatogenesis, we performed *in silico* analyses combining different methods as the first screening. The computational approach included the following steps (Supplementary Fig. S1).

Step 1, selection of protein family groups specific to plant species that produce spermatozoids. Because plant spermatozoids have structures distinct from animal sperm, such as a MLS contiguous with the spline, a spiral-shaped nucleus elongated along the spline, and multiple flagella, we hypothesized that the plant species producing spermatozoids would harbor specific genes not present in animals that are needed to form these distinctive structures. Therefore, we classified proteins of plants producing spermatozoids and animals into family groups using the OrthoFinder tool (Emms and Kelly 2019) and then selected protein family groups present only in the plants producing spermatozoids. In this step, we also used protein data forChlamydomonas, a flagellate green alga that does not produce spermatozoids, so that we could remove known flagella proteins and unrelated proteins for spermatogenesis from among the candidates by exclusion of protein families contained in Chlamydomonas. Then, 938 protein family groups remained as primary candidates (Supplementary Fig. S1, Step 1).

Step 2, extraction of protein family groups composed by genes highly expressed during spermatogenesis. We expected that genes involved in spermatogenesis should be highly expressed during spermatogenesis. Based on RNA-seq data for tissues in the vegetative and male reproductive stages in Marchantia (Higo et al. 2016) and Physcomitrella (Koshimizu et al. 2018), we extracted genes exhibiting higher expression levels during male reproductive stages compared to vegetative stages. For families consisting of multiple genes, we selected as candidates those in which all members are highly expressed at the male reproductive stages as candidates. In this step, 165 groups were retained (Supplementary Fig. S1, Step 2).

Step 3, selection of protein family groups for which member proteins exhibit low BLAST similarities with animal and Chlamydomonas proteins. In Step 1, we excluded protein family groups shared between plants and animals or Chlamydomonas. In this step, we further eliminated protein family groups that include proteins highly similar to those of animals or Chlamydomonas. Marchantia proteins in the 165 protein family groups selected in Step 2 were used for the query, and we examined sequence similarities against animals and Chlamydomonas proteins by BLASTP searching (Altschul et al. 1997; Camacho et al. 2009). For exhaustive analysis, we used ‘animals’ and ‘*Chlamydomonas reinhardtii*’ taxa of NCBI (NCBI Resource Coordinators 2018) nr datasets for the BLASTP search. When high sequence similarity to an animal or Chlamydomonas protein was detected (e-value < 0.001 or coverage > 10%), we excluded the protein family group. After this step, 31 groups remained (Supplementary Fig. S1, Step 3; and Supplementary Table S1).

Step 4, human check of the expression level data obtained in Step 2. From the Marchantia and Physcomitrella expression data used in Step 2, we selected seven genes exhibiting lower expression levels in the vegetative growth stage and substantial differences in expression levels between the vegetative and male reproductive stages. We then selected as candidates four protein family groups (Group-75, Group-89, Group-230, and Group-339) composed of seven proteins. It should be noted that although we could have initially selected genes exhibiting above-mentioned expression patterns, we preferred to use a strategy that narrows down the number of candidates after studying the functions of a broad range of proteins that could be involved in spermatogenesis (Supplementary Fig. S1, Step 4).

### Two protein family groups were selected as final candidates for functional analysis

Regarding Group-339, the function of a highly similar protein in Arabidopsis, AUG7 (AT5G17620.1), in delocalization of γ-tubulin in the mitotic spindle and phragmoplast was reported (Hotta et al. 2012). In spermatogenesis of bryophytes, centrioles serve as the microtubule organizing centers in the spindle for the final mitosis (Vaughn and Renzaglia 1998), in which γ-tubulin localizes to the centriolar centrosomes (Shimamura et al. 2004). These observations suggested that the proteins of Group-339 should play a role in γ-tubulin localization during spermatogenesis; thus, we reserved further functional analysis for Group-339. Group-75 included four proteins each of Marchantia and Physcomitrella. Because the deletion of all genes for four members would be technically difficult even in Marchantia or Physcomitrella, we also reserved this group for future analysis. The Arabidopsis protein highly similar to Group-89 is DUO3 (AT1G64570.1), which regulates male germline development, and is essential for sperm cell specification and fertilization (Brownfield et al. 2009). It would be interesting to investigate the function of this protein in spermatogenesis in bryophytes. Group-230 proteins exhibited no similarity to Arabidopsis proteins, but it did exhibit a weak similarity to Chlamydomonas BLD10 (Cre10.g418250.t1.2, CrBLD10), a cartwheel protein essential for assembly of the basal body that functions as the origin of flagella (Matsuura et al. 2004; Hiraki et al. 2007). In Phytozome ver. 5, which the analysis by Banks et al. (2011) was based on, the member of Group-230, Selaginella SELMODRAFT_427424 and PHYPADRAFT_69693 (v1.1, older model; same locus as Pp3c9_9040V3.2, but not the same exon-intron structure prediction), were placed in the same group with CrBLD10. However, the similarity was so subtle that the relation was not detected in Step 3. Current Phytozome ver. 12 “Gene Ancestry” for the viridiplantae places the Physcomitrella and Marchantia to different groups containing only mosses and Marchantia, respectively. No description on the encoding gene in Marchantia has been made, and the corresponding gene is annotated as a ‘structural maintenance of chromosomes smc family member’, with no publications reporting results of functional analyses in Physcomitrella. Thus, we decided to conduct further functional analyses for Group-89 and Group-230 consisting of one member each in Marchantia, particularly focusing our interest on spermatogenesis (Supplementary Fig. S1, Step 5).

### Mp*bld10* mutants are defective in spermatozoid formation

To analyze the roles of these two genes, we generated knock-out lines by genome editing using the CRISPR/Cas9 system (Ran et al. 2013; Sugano et al. 2018). No mutants were obtained for the *Mapoly0029s0108* gene (Group-89). For the *Mapoly0001s0460* gene (Group-230), the guide-RNA sequence was designed to target the first exon (Fig. 1A), and two independent lines harboring frameshift mutations were obtained (Fig. 1A and Supplementary Fig. S2A). Hereinafter, we refer to Group-230 family proteins as BLD10 because the phenotype and sequence analyses suggested an orthologous relationship to CrBLD10. The mutants were designated Mp*bld10-1* and Mp*bld10-2*. These mutations did not markedly affect vegetative growth of the thalli (Supplementary Fig. S2B) and formation of antheridiophores (Supplementary Fig. S2C-S2E). However, moving spermatozoids of the mutants were rarely observed for the mutants (Supplementary Movie 1-3). Intriguingly, cytoplasm elimination, nuclear elongation, and flagella formation were incomplete in the mutant spermatozoids compared with wild-type spermatozoids (Fig. 1B-1E). Immunostaining of centrin and acetylated tubulin (ac-tubulin) in spermatids was performed to observe basal bodies and the axoneme in the flagella (Fig. 1F-1I, Higo et al. 2018). Filaments of ac-tubulin were detected in a subpopulation of Mp*bld10-*spermatids, but spermatids with no detectable ac-tubulin were also observed. In spermatids positive for the ac-tubulin signal, short or coiled filaments in the cell bodies were frequently noted (Fig. 1G and 1H). The puncta signals of centrin exhibiting an abnormal size were sometimes observed in the mutant (Fig. 1I). These results suggested that Mp*BLD10* plays a crucial role in spermatozoid formation in Marchantia.

**Fig. 1.**
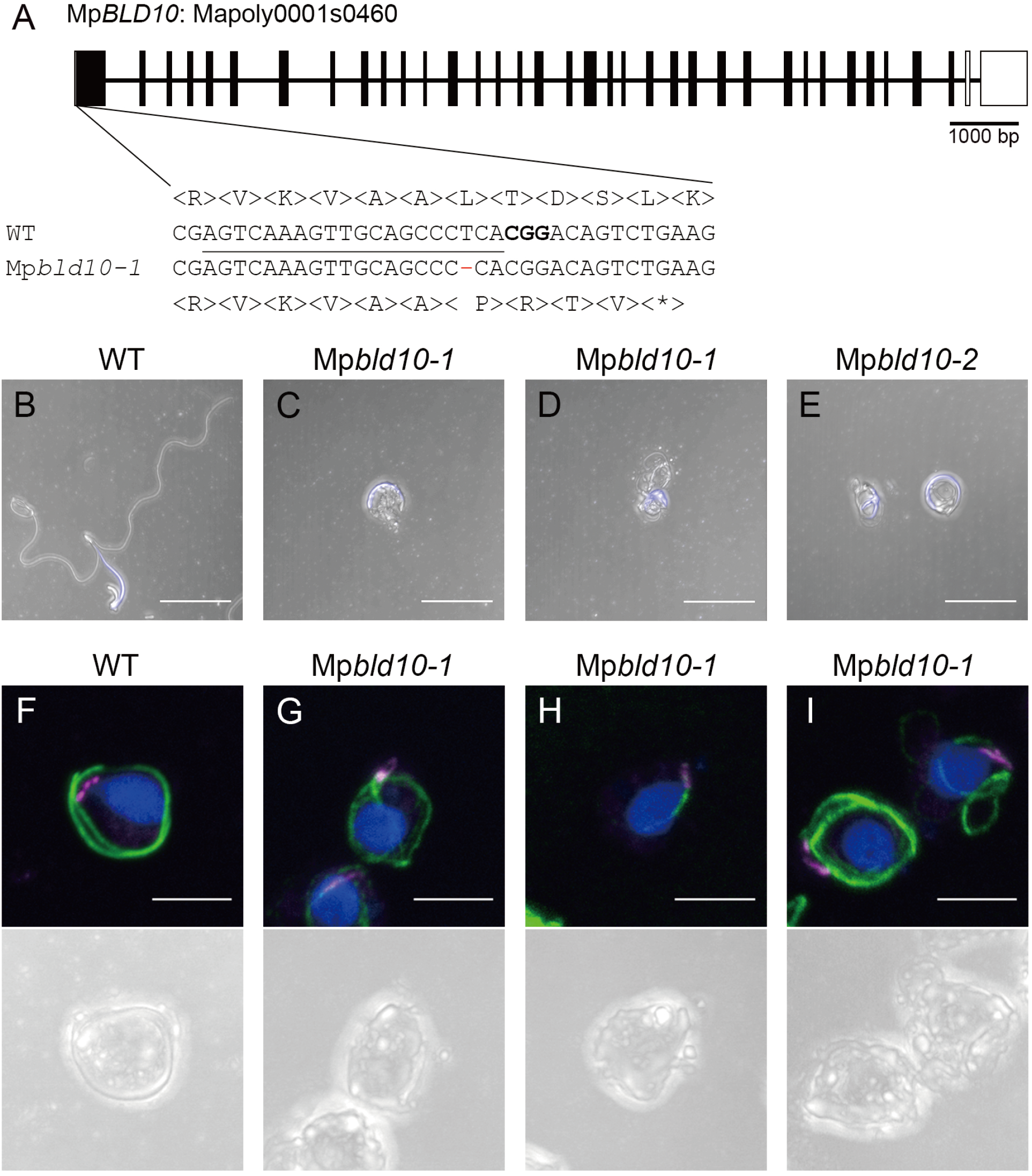
Mp*bld10* mutants exhibit severe defects in spermatozoid formation. (A) Schematic structure of the Mp*BLD10* (Mapoly0001s0460) gene. Nucleotide and amino acid sequences around mutation sites in wild-type (WT) and Mp*bld10-1* are aligned. The target and PAM sequences are indicated by underlining and bold font, respectively. (B-E) Maximum-intensity projection images of spermatozoids of wild type (B), Mp*bld10-1* (C and D), and Mp*bld10-2* (E) stained with Hoechst33342. Scale bars = 10 μm (F-I) Maximum-intensity projection images of spermatids immunostained with anti-centrin and anti-acetylated tubulin antibodies. Nuclei were visualized using Hoechst33342. Blue, green, and magenta pseudo colors indicate Hoechst33342, Alexa 488, and Alexa 594, respectively. Scale bars = 5 μm.

To examine the effect of the mutation in Mp*BLD10* at an ultrastructural level, we conducted a transmission electron microscopy (TEM) analysis of spermatids and spermatozoids of the Mp*bld10-1 mutant*. The flagella of wild-type spermatids contained an axoneme comprising two central microtubules and surrounding nine doublet microtubules (Fig. 2A). In the Mp*bld10-1 mutant*, however, a major population of spermatids did not harbor flagella, and the flagella formed in a subpopulation of spermatids exhibited a disordered axoneme structure (Fig. 2E). No structural abnormalities were detected in the Mp*bld10-1* mutant MLS, a structure unique to plants that is attached to the anterior mitochondrion in spermatids and consists of the spline, which is the uppermost stratum containing arrayed microtubules, and a lower strata with high electron densities, namely, the lamellar strip (Fig. 2B and 2F). We also observed wild-type basal bodies, which contain nine triplet microtubules and are attached to the spline of the MLS (Fig 2C and 2D). In the Mp*bld10-1 mutant*, the wild type-like basal bodies were only occasionally observed, and amorphous electron-dense regions were frequently observed instead of basal bodies (Fig. 2G and 2H). These results strongly suggested that Mp*BLD10* is required for correct assembly of the basal body and flagella during spermatogenesis. Furthermore, we observed that the Mp*bld10-1* spermatozoids also exhibited a defect in chromatin compaction in the nucleus (Fig. 2I and 2J). Thus, Mp*BLD10* might also be involved in chromatin organization during spermiogenesis in Marchantia.

**Fig. 2.**
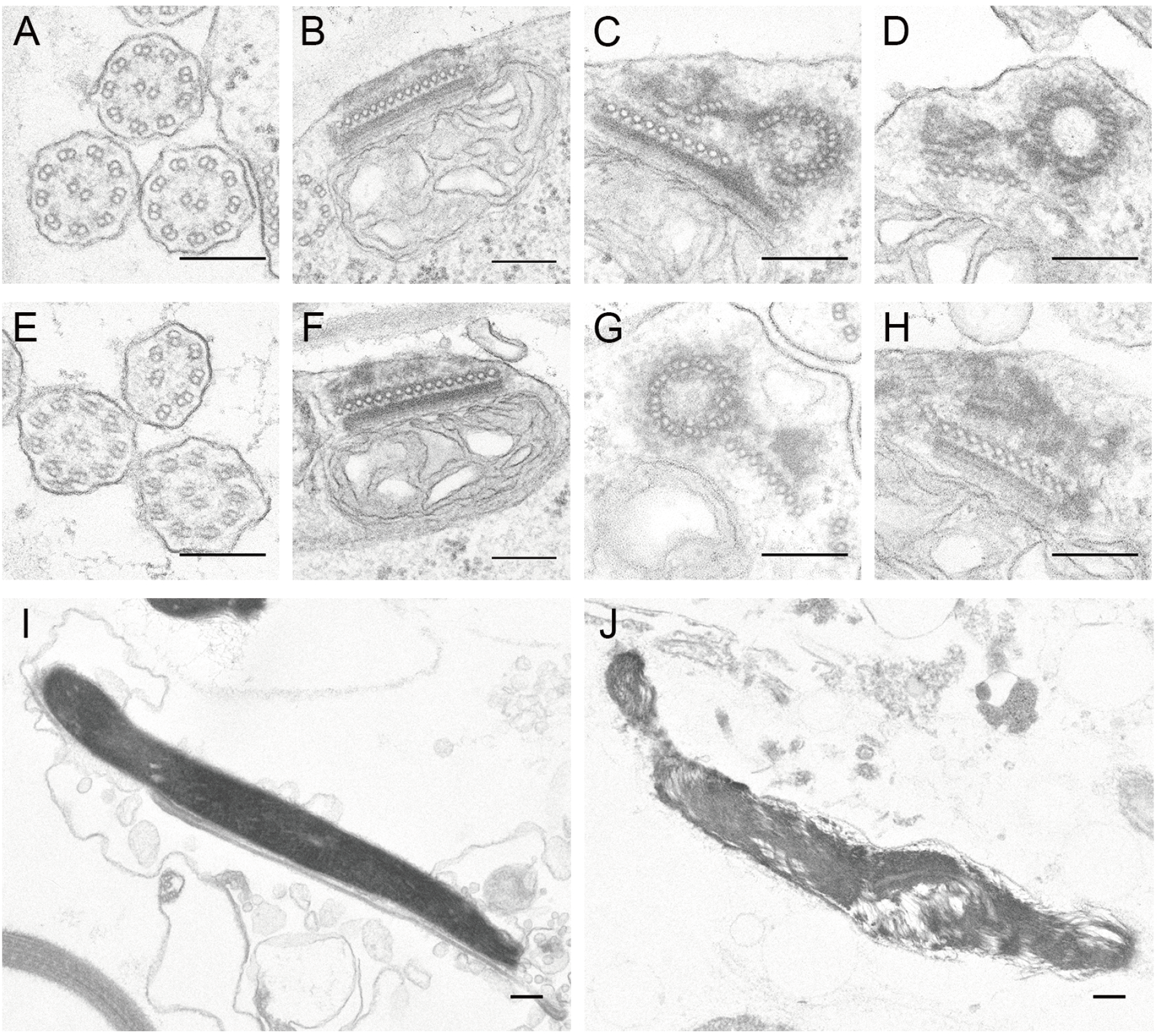
Transmission electron microscopy (TEM) of spermatids and spermatozoids in wild type and Mp*bld10-1*. (A-H) TEM images in spermatids of wild type (A-D) and Mp*bld10-1* (E-H). Axonemes in flagella (A and E), multilayered structures (B and F), and basal bodies (C, D, G, and H) are shown. (I and J) TEM images of nuclei in spermatozoids of wild type (I) and Mp*bld10-1* (J). Scale bars = 200 nm.

### PpBLD10 is required for formation of basal bodies and flagella

To further examine the functions of the Group-230 proteins (BLD10s) in bryophytes, we analyzed the ortholog of Mp*BLD10* in Physcomitrella, the model of moss readily amenable to gene targeting and genome editing. The Group-230 in Physcomitrella consists of only *Pp3c9_9040V3*.*2* (Pp*BLD10*). We generated knock-out mutants of this gene using the CRISPR-Cas9 system. Pp*bld10-22*, which has an 11-bp deletion in the first exon, and Pp*bld10-30*, which has an approximately 10-kbp deletion between exon 1 and exon 29 of the transcript XM_024529611.1, were used in further analyses (Supplementary Fig. S3). Although these mutants did not exhibit any marked defects in protonemata, gametophores, and gametangia (Supplementary Fig. S4), spermatozoids of these mutants were not motile; no moving mutant spermatozoids were observed (Supplementary Movie 4-6). Spermatids lacking the signal or with coiled filaments were observed in the mutants immunostained for ac-tubulin (Supplementary Fig. S5C-S5F), indicating that the mutants are defective in flagella formation, similar to Marchantia. No marked defects in elimination of the cytoplasm and nuclear elongation of the mutants were observed (Supplementary Fig. S6A-S6C).

In the TEM analysis of spermatids and spermatozoids, no axoneme structure was observed in most of the spermatids in Pp*bld10-30* (Supplementary Fig. S7A and S7C), and amorphous electron-dense regions were observed instead of basal bodies (Supplementary Fig. S7B and S7D). With regard to the MLS, no marked defects were observed in the mutant, similar to the Mp*bld10-1* mutant (Supplementary Fig. S7B and S7D). In contrast to the Mp*bld10-1*, no defect in chromatin compaction in the nucleus was observed in Pp*bld10-30* (Supplementary Fig. S7E and S7F). These results suggest that PpBLD10 functions in basal body- and flagella assembly, but the requirement for PpBLD10 in chromatin compaction and nuclear formation differs from the case in Marchantia.

### MpBLD10 and PpBLD10 localize in basal bodies during flagella formation

We next examined the subcellular localization of MpBLD10. We generated a transgenic line of Marchantia expressing mCitrine-fused MpBLD10 driven by its own promoter in Mp*bld10-1*. Expression of mCitrine-MpBLD10 restored the defects in spermatozoid formation and motility in the mutant, indicating that this chimeric protein retains the authentic function (Supplementary Movie 7). In this transgenic line, we traced spermiogenesis according to developmental stage as previously defined (Minamino et al. 2021). In stage 0 spermatids, mCitrine-MpBLD10 was observed as two closely aligned rod-like structures (Fig. 3A). In stage 1 spermatids, mCitrine-positive structures were located at the base of the flagella (Fig. 3B). In stage 2, the mCitrine-MpBLD10 signal became weak, then ultimately disappeared in subsequent stages (Fig. 3C-3F). To verify the nature of these structures, we performed co-immunostaining with centrin and ac-tubulin. As shown in Fig. 3G, mCitrine-MpBLD10 was localized in close association with centrin at the proximal side of the flagella (Fig. 3G). These results suggest that MpBLD10 localize in the basal body during flagella formation and is then degraded after flagella formation.

**Fig. 3.**
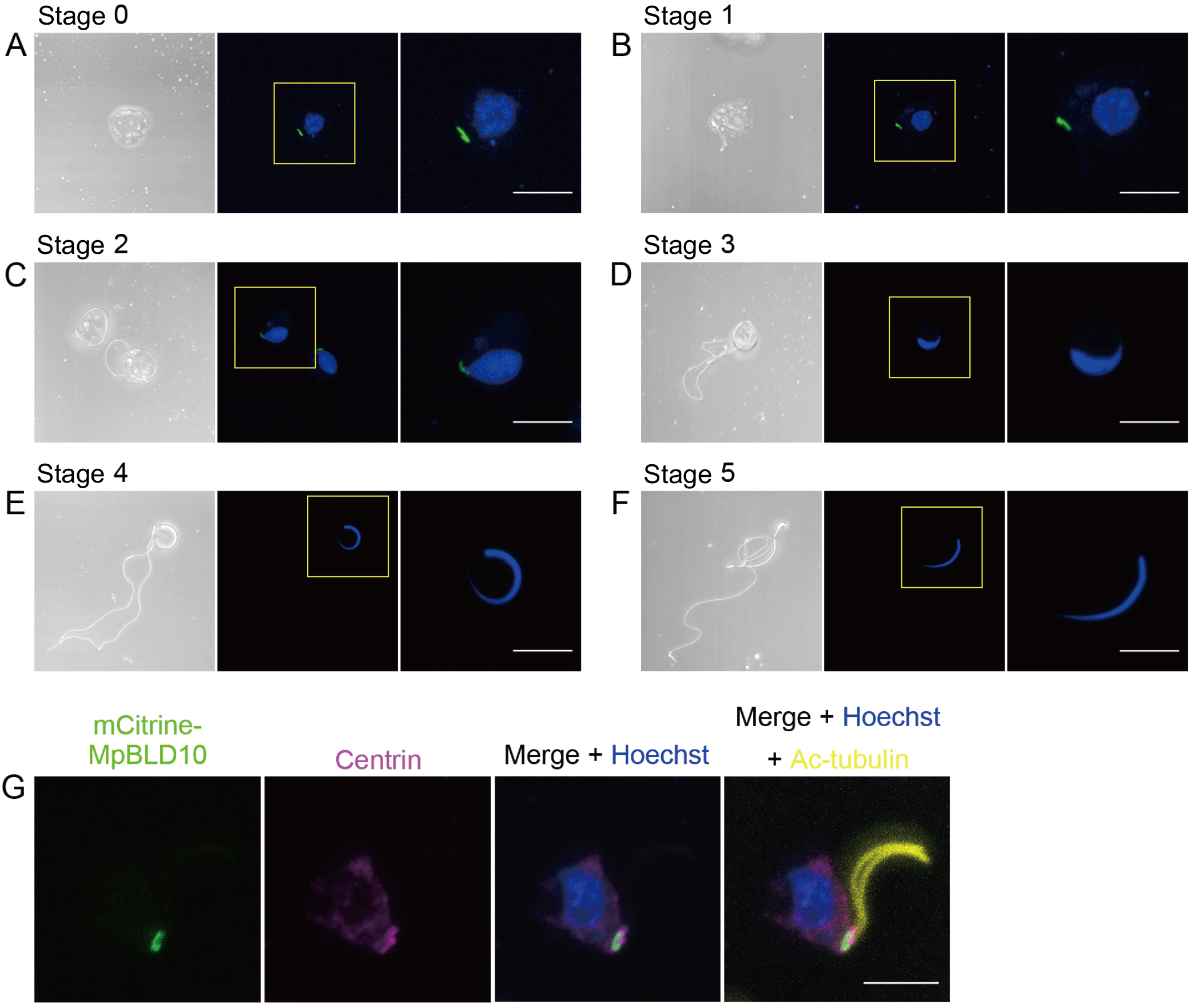
Subcellular localization of the MpBLD10 protein. (A-F) Differential interference contrast microscopy (DIC) and maximum-intensity projection images of spermatids and spermatozoids expressing mCitrine-MpBLD10 (green) driven by its own promoter at stage 0 (A), stage 1 (B), stage 2 (C), stage 3 (D), stage 4 (E), and stage 5 (F); nuclei were stained with Hoechst33342 (blue). Developmental stages were classified according to Minamino et al. (2021). (G) Maximum-intensity projection images of a spermatid expressing mCitrine-MpBLD10 (green) immunostained with anti-centrin (magenta) and anti-acetylated tubulin (yellow) antibodies. The nucleus was stained with Hoechst33342 (blue). Scale bars = 5 μm.

The localization of PpBLD10 in Physcomitrella was examined using Citrine knock-in lines (Supplementary Fig. S3). Spermiogenesis was observed according to the developmental stages defined for Marchantia (Minamino et al. 2021) (Supplementary Fig. S8A-S8E), but in Physcomitrella, flagella formation was slower than in Marchantia and began when the nuclear shape was spindle-like; thus, stages 0 and 1 as defined for Marchantia are indistinguishable in Physcomitrella. PpBLD10-Citrine signals were observed in stage 0-1 and stage 2 but disappeared in subsequent stages, as in Marchantia (Supplementary Fig. S8A-S8E). In stage 0-1, the PpBLD10-Citrine was observed as puncta (Supplementary Fig. S8A); in stage 2, the signal was detected in a wider region of the basal part of the flagella as compared with stage 0-1 (Supplementary Fig. S8B and S8F). The positional relationship between PpBLD10-Citrine and centrin in Physcomitrella could not be observed because the anti-centrin antibody did not immunostain Physcomitrella spermatids (data not shown).

### BLD10s exist in streptophytes with flagella and evolved fast in land plants

CrBLD10, to which MpBLD10 and PpBLD10 exhibit weak similarity, is an ortholog of *Homo sapiens* CEP135 (HsCEP135) (Carvalho-Santos et al. 2010), which plays a role in early basal body/centriole biogenesis (Kleylein-Sohn et al. 2007). To obtain information on BLD10 proteins in the streptophyte lineage and to assess their orthology to the BLD10/CEP135 family, we searched the genome, transcript, and protein sequences of species ranging from streptophyte algae to angiosperms (angiosperms: *Arabidopsis thaliana* and *Oryza sativa*; gnetophytes: *Gnetum montanum*; Pinaceae: *Picea abies* and *Pinus taeda*; Ginkgo: *Ginkgo biloba*; monilophytes: *Salvinia cucullata* and *Azolla filiculoides*; lycophytes: *Selaginella moellendorffii*; bryophytes: *Anthoceros punctatus*; Zygnematophyceae: *Penium margaritaceum, Mesotaenium endlicherianum*, and *Spirogloea muscicola*; Coleochaetophyceae: *Coleochaete orbicularis*; Charophyceae: *Chara braunii*; Klebsormidiophyceae: *Klebsormidium nitens*; Mesostigmatophyceae: *Mesostigma viride*; Chorokybophyceae: *Chlorokybus atmophyticus*). In Arabidopsis, Oryza, Gnetum, Picea, Pinus, Penium, Mesotaenium, and Spirogloea, no hit sequences were obtained. Some of the hit sequences were reconstructed based on RNA-seq data and similarity to preliminary alignments. Finally, Ginkgo (GbBLD10a and GbBLD10b), Salvinia (Sacu_v1.1_s0007.g003760), Azolla (AfBLD), Selaginella (SmBLD10), Anthoceros (Apun_evm.model.utg000038l.487.1), Coleochaete (GBSL01053926.1), Chara (CbBLD10), Klebsormidium (kfl00353_0080_v1.1), Mesostigma (MvBLD10), and Chlorokybus (Chrsp112S01623) were subjected to the alignment and phylogenetic analyses together with MpBLD10, PpBLD10, and CrBLD10 and non-plant BLD10/CEP135 family sequences from *H. sapiens* (HsCEP135), *Drosophila melanogaster* (DmBLD10), and *Tetrahymena thermophila* (TtBLD10) as outgroup sequences (Fig. 4A and Supplementary Fig. S9). Additionally, we included the additional sequences *Adiantum capillus-veneris* MBC9850943.1, *Chlamydomonas eustigma* CEUSTIGMA_g448.t1, and *Bombus impatiens* XP_012244165.1 to stabilize the alignment and phylogenetic tree by dividing long branches. Here, among multiple potential isoforms, ref_seq protein XP_024385379.1 was used for the phylogenetic analysis because this isoform of PpBLD10 was well supported by RNA-seq data and fit the alignment better.

**Fig. 4.**
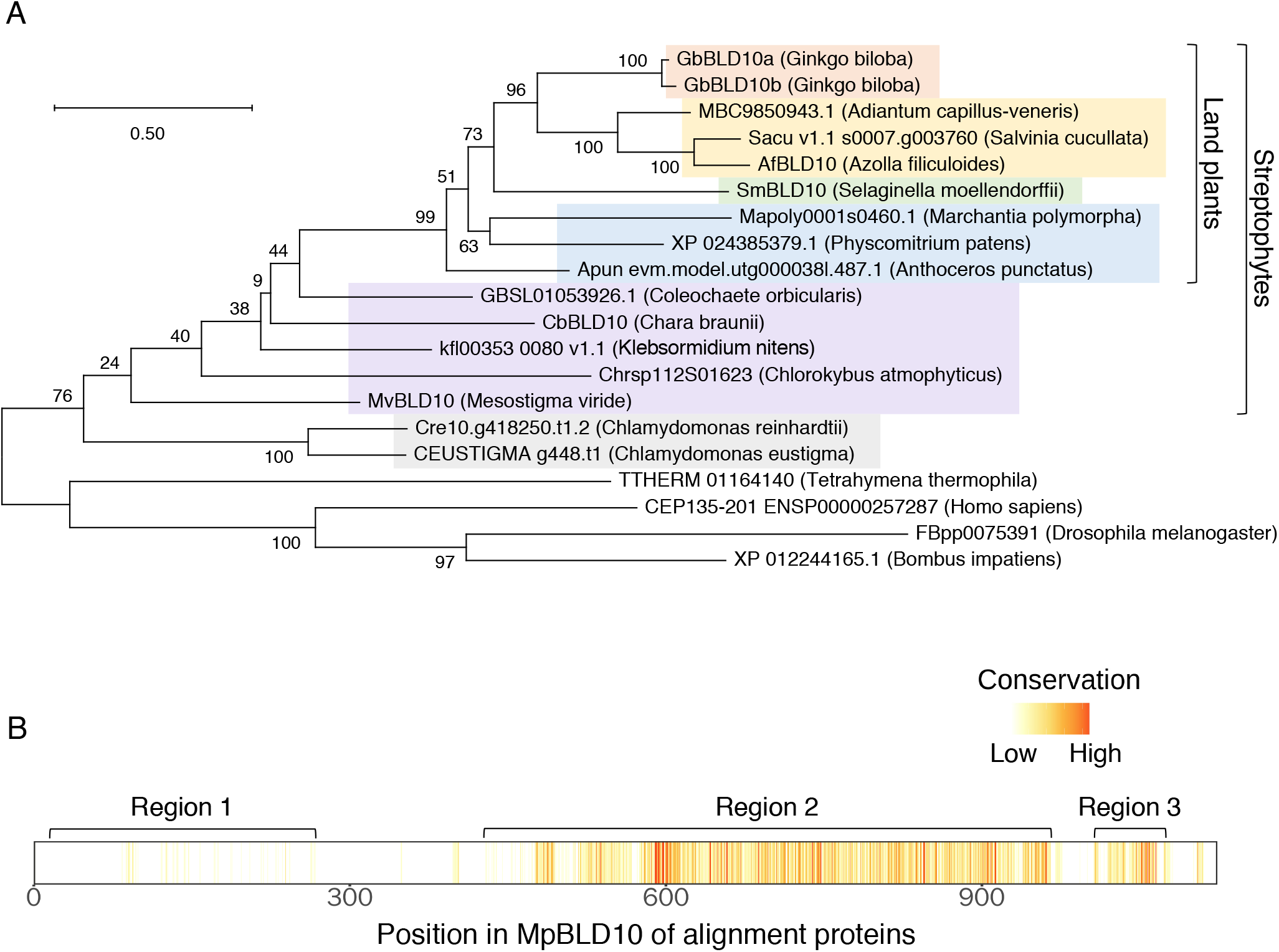
Comparisons of BLD10/CEP135 family protein sequences. (A) Phylogenetic tree of BLD10/CEP135 family proteins. Out-groups are Tetrahymena, human, and insects. Salmon, yellow, green, blue, purple, and gray background colors indicate gymnosperms, monilophytes, lycophytes, bryophytes, streptophyte algae, and chlorophytes, respectively. Branch lengths are proportional to the estimated number of amino acid substitutions/site (scale upper left). (B) Architecture of the MpBLD10 sequence (gaps removed) with conservation level among BLD10 proteins in the streptophytes in (A). Three conserved regions are indicated. Values of conservation levels were obtained using Jalview software.

In the phylogenetic analysis, the green plant, streptophyte, and land plant genes formed a clade (Fig. 4A). Among land plants, the setaphytes (mosses + liverworts), and euphyllophytes formed a clade with low and high bootstrap supports, respectively. Outside of land plants, Coleochaetophyceae, Charophyceae, Klebsormidiophyceae, and Chlorokybophyceae branched in that order, with low bootstrap support. The branches of the land plants were longer than those of streptophyte algae (Fig. 4A). The molecular clock hypothesis was rejected between land plants and streptophyte algae, with Chlamydomonas as out-group by Tajima’s relative rate test (p < 0.05; Tajima 1993), indicating an increase in the evolutionary rate of BLD10s in the land plant lineage. Upon application of a local clock model in PAML (Yang 2007) to the dataset for green plants, such that the branches of land plants after divergence from Coleochaetophyceae has different rates than all other branches, land plants had an approximately 2-fold higher evolutionary rate than green algae.

### Green plant BLD10s have a novel conserved domain close to the C-terminus

**Functional regions were reported for** CrBLD10 and HsCEP135: a probably essential region for the function of CrBLD10 (residues 850 - 1,050; Hiraki et al. 2007), and three binding regions in HsCEP135 (microtubule-binding region, residues 1 - 190; CPAP-binding region, residues 50 - 460; hSAS-6-binding region, residues 416 - 1,140; Lin et al. 2013). Among streptophyte BLD10s, three conserved regions were identified (Fig. 4B and Supplementary Fig. S9). Region 1 does not seem to exist in Selaginella, because the N-terminal region is annotated as a separate protein in the ref_seq annotation (XP_024518287.1). The N-terminus of region 1 was not detected in either Ginkgo or Adiantum. As such, the level of conservation shown in Fig. 4B and Supplementary Fig. S9 is low, but region 1 is conserved in other streptophytes. The microtubule-binding region and the N-terminal half of the CPAP-binding region correspond to region 1 in streptophytes, but a large proportion of the BLD10/CEP135 family is so divergent that the conserved residues are rare (Supplementary Fig. S9). The C-terminal half of the CPAP-binding region was found to be mostly conserved among green algae including Coleochaete, but it was shortened in land plants. Region 2 includes the probably essential region of CrBLD10 and overlaps with the hSAS-6-binding region. Region 2 is a long and conserved, but a large deletion of 49 residues in the CrBLD10 essential region was found in MpBLD10 (Supplementary Fig. S9). A review of RNA-seq data mapped to the region confirmed that the loss in MpBLD10 was not due to an annotation error skipping an exon. The 27 N-terminal residues are highly conserved in green plants. In streptophytes, region 3 is a 65-residues long highly conserved region close to the C-terminus (Supplementary Fig. S9). Although some residues aligned, the distances to human, fly, and Tetrahymena were so large that the homology is obscure, and the alignment was unstable to additional insect sequences. Similarity to outside of land plants cannot be detected through PSI-BLAST (Altschul et al. 1997) with land plant sequences in NCBI, and no conserved domain was found in the conserved domain database (CDD; Lu et al. 2020). An examination of the alignment revealed that, CrBLD10 has region 3 starting with the signature (KR)XX(ED)LE and extending to (LVM)(LV)X(LI)(LM)(SA)(KR)(VL)(DE)X(DE)(RK) except Q rich block, thus it was judged to be homologous. Region 3 constitutes a novel conserved domain in green plants. A 3-amino acid (aa) deletion and lower conservation in the N-terminus of the region was noted in Klebsormidium. The intron-exon structure is fully supported by RNA-seq data.

## Discussion

In this study, we established a pipeline that allows for efficient and rapid searches of protein family groups involved in plant spermatogenesis through a computational analysis of large omics datasets stored in publicly available online databases. Our *in silico* approach involves two primary steps: (i) selection of protein families specific to streptophytes that produce spermatozoid, and (ii) selection of protein families encoded by genes highly and predominantly expressed during spermatogenesis. Using this pipeline, we extracted seven genes from approximately 19,000 Marchantia genes belonging to four protein family groups (Group-75, Group-89, Group-230, and Group-339). Among these protein families, we conducted functional analyses of Group-89 and Group-230 in Marchantia. No Group-89 mutants were obtained, but we successfully generated loss-of-function mutants for Group-230 (BLD10s). We then found that the BLD10s play a crucial role in spermatogenesis in Marchantia and Physcomitrella, thus showing the effectiveness and accuracy of our computational selection approach. This method is applicable to the identification of genes involved in a variety of biological processes other than spermatogenesis using large omics datasets.

Most spermatozoids of the Mpbld10 mutants did not have flagella. A few spermatozoids possessed short or coiled filaments within the cytoplasm (fig. 1G-1I), likely corresponding to incomplete flagella observed in a TEM image (fig. 2E). Similar results were observed in Pp*bld10* mutants (Supplementary Fig. S5C-S5F and Supplementary Fig. S7C). Generally, flagella are observed outside the cell body in the early stages of spermatogenesis (Minamino et al. 2021), but the immature flagella-like structures remained inside the cell in the Mp*bld10* and Pp*bld10* mutants. In addition to this shared characteristic of Marchantia and Physcomitrella, the Mp*bld10* mutants also exhibited defective cytoplasm elimination, chromatin compaction, and nuclear elongation (Fig. 1C-1E and Fig. 2J), which were not observed in the Pp*bld10* mutants (Supplementary Fig. S6B and S6C and Supplementary Fig. S7F). MpBLD10 and PpBLD10 were localized at the base of forming flagella in the early stages of spermiogenesis and disappeared after flagella formation was completed (Fig. 3A-3F and Supplementary Fig. S8A-S8E). This localization, together with the close association between MpBLD10 and centrin at the base of the flagella (Fig. 3G), suggests that MpBLD10 and PpBLD10 are basal body proteins that function in basal body assembly. MpBLD10 is likely also involved in chromatin reorganization during spermiogenesis. The additional effect of the Mp*bld10* mutation on phenotypes shared between Mp*bld10 a*nd Pp*bld10* suggests that MpBLD10 plays an additional role during spermiogenesis and that the functions of MpBLD10 and PpBLD10 partially diverged during bryophyte evolution.

In the phylogenetic analysis, the branching order among streptophyte algae was congruent with an organism tree based on phylotranscriptomic analyses (Wickett et al. 2014; Puttick et al. 2018; Leebens-Mack et al. 2019) with low bootstrap support (Fig. 4A). Although the monophyly of land plant genes was supported by a high bootstrap value, among land plants, the tree was congruent with the phylogeny of organisms with low bootstrap supports, except for the position of the hornwort gene. The placement of hornworts sister to all other land plants has been observed at a low frequency in low copy number gene phylogenies of land plants (Li et al. 2020). Based on the congruence to the phylogeny, these streptophyte BLD10s are putative orthologs of CrBLD10 (fig. 4A). Flagellate species in streptophytes have BLD10; conversely, species without flagella (i.e., Zygnematophyceae, conifers/gnetophytes, and angiosperms) lost BLD10 (Fig. 5). BLD10/CEP135 family proteins function in assembly of the basal body/centriole, which is involved in cell division and serves as the basis of flagella/cilia in human (Kleylein-Sohn et al. 2007; Lin et al. 2013), fly (Mottier-Pavie and Megraw 2009; Carvalho-Santos et al. 2012), Tetrahymena (Bayless et al. 2012), and Chlamydomonas (Matsuura et al. 2004; Hiraki et al. 2007). MpBLD10 and PpBLD10 also play a role in basal body assembly, similar to other BLD10/CEP135 family proteins. In accordance with the differences in BLD10 sequences between land plants and Chlamydomonas, the mutant phenotypes differed from the mutants of other BLD10/CEP135 family proteins; basal bodies were completely lacking and flagella never observed in Cr*bld10*, but incomplete basal bodies and flagella were formed in the Mp*bld10* and Pp*bld10* mutants. Although DmBLD10 remains after the development of sperms in fly (Mottier-Pavie and Megraw 2009), MpBLD10 and PpBLD10 disappeared, suggesting that MpBLD10 and PpBLD10 are necessary only during flagella formation, and no longer needed after the formation of flagella. In addition, Mp*bld10* mutants exhibited defects in cytoplasmic reduction and nuclear elongation during spermiogenesis, and this phenotype was observed specifically in Marchantia but not in Physcomitrella. Changes in the role of basal body/centriole in the life cycle or environment of fertilization (free water, limited water, or pollination droplet) in land plants might have affected the evolutionary rate of BLD10s in land plants through positive selection or a relaxation of purifying selection, which cannot be discerned from the current data. Note that centrioles are not observed during cell division in land plants (Buschmann and Zachgo 2016) (Fig. 5). Despite *Klebsormidium nitens* NIES-2285 do not show flagellated cells under laboratory conditions, BLD10 expression is detected in RNA-seq data, implying their role in cell division rather than flagella formation. BLD10 probably played a dual role in the flagella formation and cell division in the ancestral green algae and then lost the role in cell division in land plants after other microtubule organization mechanisms for chromosome separation during cell division were established; thus, the protein has a sperm-specific function in bryophytes, lycophytes, and ferns. BLD10 gene was lost entirely in conifers/gnetophytes and angiosperms, consistent with the loss of flagellated sperm cells.

**Fig. 5.**
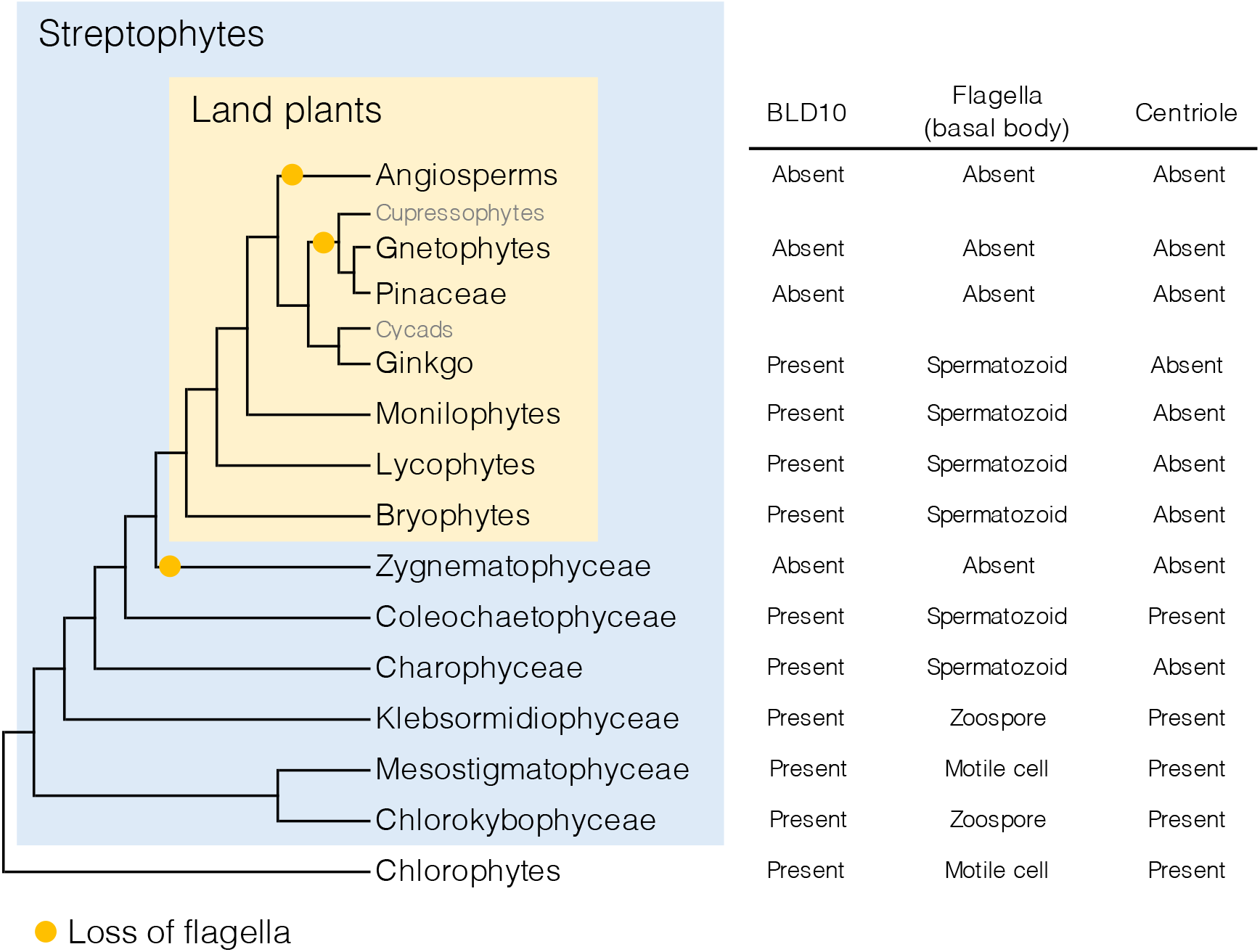
Existence of BLD10 proteins, flagella (basal body), and centriole during cell division in the plant species. Orange dots show presumed flagella loss. Groups indicated by gray color were not investigated due to insufficient sequence data. For phylogenetic relationships and presence of centrioles, refer to Puttick et al. (2018) and Buschmann and Zachgo (2016), respectively.

We successfully identified genes that function in plant spermatogenesis using an *in silico* analysis. Two identified proteins, MpBLD10 and PpBLD10, are basal body proteins that exhibit a higher evolutionary rate despite the importance of their role in assembly of the basal body during spermatogenesis. MpBLD10 possibly acquired a new role in spermatozoids formation—specifically, in cytoplasm elimination and nuclear elongation—through plant evolution. Notably, the relationship that streptophyte BLD10s are putative orthologs of BLD10 was not found in a simple run of OrthoFinder, and identified only after a more detailed analysis. The importance of gene loss in the evolution was recently documented in various fields/cases, and phylogenetic distribution of the gene involved in a particular trait coincides with the distribution of the trait (Glastad et al. 2011; Zhang et al. 2013; Lin et al. 2017; Griesmann et al. 2018; Sharma et al. 2018; Gluck-Thaler et al. 2020). In this analysis, we demonstrated that the process can be reversed; that is, a gene responsible for the trait can be identified by searching for a gene whose phylogenetic distribution coincides with the trait combined with the expression at the site the trait is observed. The results of this study thus highlight the power of combinatorial analyses with expression and multiple independent losses of traits and underlying genes during evolution.

## Materials and Methods

### *In silico* screening

Ortholog groups were predicted using OrthoFinder (v2.3.3, Emms and Kelly 2019) with the protein sequences of the species shown in Supplementary Table S2. For expression level quantification in Marchantia and Physcomitrella, transcripts per million (TPM) values were calculated using RSEM (v1.3.1, Li et al. 2011) with Bowtie2 (v2.3.5.1, Langmead et al. 2012) for RNA-seq data listed in Supplementary Table S3 using the reference transcriptome datasets listed in Supplementary Table S2.

### Sequence searching and reconstruction of gene models

BLD10 proteins in the streptophytes were searched against the genome, transcript, and protein sequences using BLAST (v2.10.1+, Altschul et al. 1997; Camacho et al. 2009). In the protein datasets, BLASTP searching was performed using MpBLD10 against the species shown in Supplementary Table S2. Hit protein sequences are listed in Supplementary Table S4. The lengths of hit sequences in Ginkgo and Chara were <600 aa (full length of MpBLD10 is 1,123 aa). Because the close gene ids implied close positions in the genome, genomic locations were investigated. Gb_13822 is at the 279 Mb position on Chr12; Gb_03087-Gb03089, Gb_30854, and Gb_30855 are close to the 630 Mb position on Chr12; Gb_39501 is close to the 277 Mb position on Chr7. Two gene models on Chr12 were reconstructed based on mapping of RNAseq data (*GbBLD10a* and *GbBLD10b*), and no reads were found for Gb_39501. In Chara, we constructed a presumptive transcript (*CbBLD10*) from the genome and RNA-seq data (Nishiyama et al. 2018). Azolla sequence Azfi_s0013.g013382 had a deletion in the C-terminal region. The missed exons were supplied based on the RNA-seq data (Li et al. 2018), and a gene model (Af*BLD10*) was constructed. In Selaginella, the hit sequence SELMODRAFT_427424 had three deletions and one insertion relative to most of the green plant BLD10s in the initial alignment. By investigating the genomic sequence with the preliminary alignment as a guide, a gene model more similar to the conserved consensus was constructed (*SmBLD10*), though the RNA-seq data of sperm-producing tissue in Selaginella were insufficient. Although SELMODRAFT_427424 resides on scaffold_90, another copy (allele) is present on scaffold_104, but contained an assembly gap (stretch of Ns) and was not annotated. Further, transcriptome shotgun assemblies (TSA) were searched using TBLASTN at NCBI (NCBI Resource Coordinators 2018) with CrBLD10. Hit transcript sequences for *Mesostigma viride* and *Coleochaete orbicularis* were obtained (Supplementary Table S4). Mesostigma had split hits of the different contigs with reasonable similarity, presumably constituting the corresponding protein. The three contigs were mapped to scaffold_80 between 197,517 and 234,765. Thus, a gene model encoding 1744 residues (MvBLD10) was constructed for the region mostly based on RNA-seq mapping and a little guess work. The *Coleochaete orbicularis* contig appeared to contain a complete coding sequence. The reconstructed gene models are shown in Supplementary Data 1 and 2.

### Method for reconstructing gene models based on RNA-seq data or alignment similarity

Ginkgo, Azolla, Selaginella, Chara, and Mesostigma BLD10 sequences were reconstructed according to the following method. RNA-seq data were previously published or downloaded from SRA using fastq-dump (Supplementary Table S3). The RNA-seq data were mapped to a corresponding reference genome using hisat2 (v2.2.1; Kim et al. 2019). The mapped bam files were sorted according to the coordinates using samtools (Danecek et al. 2021) and then indexed. The reference sequence and annotations in gff files were loaded to JBrowse-1.15.4 (Buels et al. 2016) using prepare-reference and flatfile-to-json, respectively. The indexed bam files were loaded using add-bam-track. After preparing the data directory of JBrowse, the data were connected to Apollo-2.1.0 for manual editing. The gene model or bam read alignment was chosen for the new gene models and merged or extended in the user model editing pane. For the *Ginkgo biloba* reference, the bam file could not be indexed due to its large reference size (>512 Mb). Thus, the reference was cut to 500 Mb, and Chr12, at >500 Mb, were named as ‘Chr12b’ for subsequent processing. The records of bam files were screened for Chr7 at <500 Mb and Chr12, and records for Chr12 >500 Mb were placed to Chr12b, and positions were subtracted by 500 million. The reference genome gff was similarly edited. Thus, Chr7, Chr12 up to 500 Mb, and Chr12 over 500 Mb were loaded to JBrowse and apollo2.

### Phylogenetic analyses

The protein sequences HsCEP135 (ENSP00000257287), DmBLD10 (FBpp0075391), and TtBLD10 (TTHERM_01164140) were obtained via Ensembl (Yates et al. 2020), FlyBase (Larkin et al. 2020), and TGD (Stover et al. 2006), respectively. *Adiantum capillus-veneris* MBC9850943.1, *Chlamydomonas eustigma* CEUSTIGMA_g448.t1, and *Bombus impatiens* XP_012244165.1 were obtained from NCBI. Multiple sequence alignment was performed using MAFFT (Katoh and Standley 2013) with the accurate option E-INS-i in Jalview software (v2.10.5, Waterhouse et al. 2009). The conservation level was calculated based on that used in the AMAS method for multiple sequence alignment analysis (Livingstone and Barton 1993) in Jalview. A phylogenetic tree was constructed based on the multiple alignment with complete deletion of gap sites using the maximum-likelihood method of MEGA X software (Kumar et al. 2018), with 1,000 bootstrap replicates. Jones-Taylor-Thornton (JTT) matrix-based model (Jones et al. 1992) with gamma-distribution among sites and subtree-pruning-regrafting - extensive (SPR level 5) were used for amino acid substitution model and heuristic methods. Protein regions used in the analyses are shown in Supplementary Fig. S9 (orange boxes). Differences in branch length between land plants and green algae were compared using the local clock model in PAML4 (v4.9j; Yang 2007). Branches in land plants and the branch leading to land plants after divergence with Coleochaetophyceae were assigned category #1 in the comparison. JTT model was used for amino acid rate. Other parameters were the same as those of the ‘aaml.ctl’ file distributed with the software.

### Plant materials

Male accession of Marchantia, Takaragaike-1 (Tak-1), was used for observing spermatogenesis in this study. Plants were grown on 1/2× Gamborg’s B5 medium containing 1.4% (w/v) agar at 22°C in continuous white light. Induction of sexual organ generation by far-red irradiation was performed as described previously (Chiyoda et al. 2008).

Physcomitrella Gransden ‘Cove-NIBB’ line (Nishiyama et al. 2000) was used as the wild type. Physcomitrella was cultured on BCDAT medium with 0.8% (w/v) agar at 25 °C under continuous light conditions for protonemata (Nishiyama et al. 2000). Protonemata were transplanted into sterile peat pots (Jiffy-7; Jiffy Products International AS) and cultured approximately one month at 25 °C in continuous light for gametophores. To obtain gametangia and sporophytes, the peat pots containing gametophores were incubated at 15 °C under short-day (8-h light and 16-h dark) conditions (Sakakibara et al. 2008).

### Transformation of Marchantia

To construct vectors for genome editing, target sequences were selected using CRISPR direct (https://crispr.dbcls.jp/) (Naito et al. 2015), and double-stranded oligonucleotides of the target sequences were inserted into the pMpGE_En03 vector (Sugano et al. 2018). The resultant gRNA cassettes were introduced into the pMpGE010 vector using Gateway LR Clonase II Enzyme Mix (Thermo Fisher Scientific). To express Citrine under regulation of the Mp*BLD10* promoter, 5 kb of the 5’ flanking region of Mp*BLD10* with a SmaI site was introduced into the pENTR/D-TOPO vector (Invitrogen). The chimeric sequence was introduced into pMpGWB307 (Ishizaki et al. 2015) using Gateway LR Clonase II Enzyme Mix. To construct the mCitrine-MpBLD10 vector, a genomic fragment containing the coding region, intron, and 4 kb of the 3’ flanking region of Mp*BLD10* was inserted into the SmaI site of the pENTR/D-TOPO vector containing _*pro*_Mp*BLD10* by using an In-Fusion HD cloning kit (Clontech). A silent mutation was introduced into the PAM site by inverse PCR. The CDS of monomeric Citrine was introduced into the SmaI site of the pENTR/D-TOPO vector containing the Mp*BLD10* genomic fragment by using an In-Fusion HD cloning kit (Clontech). The chimeric sequence was introduced into pMpGWB301 (Ishizaki et al. 2015) using Gateway LR Clonase II Enzyme Mix. The primer sequences are listed in Supplementary Table S5.

Transformation of Marchantia was performed according to previously described methods (Ishizaki et al. 2008; Kubota et al. 2013). Transgenic lines were selected with 10 mg l^-1^ hygromycin B and 100 mg l^-1^ cefotaxime for pMpGE010, and 0.5 μM chlorsulfuron and 100 mg l^-1^ cefotaxime for pMpGWB301.

### Transformation of Physcomitrella

To perform CRISPR-mediated mutagenesis, we designed oligodeoxynucleotides used as single-guide RNAs (sgRNAs) targeting Pp*BLD10* (Supplementary Fig. S3B) using CRISPRdirect (https://crispr.dbcls.jp/; last visited April, 2021). The annealed oligodeoxynucleotides were cloned into the BsaI site of pPpU6-sgRNA (LC494193) (Gu et al. 2020). To generate genome editing mutants, protoplasts were co-transformed with a total of 6 µg of circular DNAs divided as follows: 2 μg of pPpU6-Pp*BLD10*-sgRNA#1 or pPpU6-Pp*BLD10*-sgRNA#2, 2 μg of pAct-Cas9 (Collonnier et al. 2017), and 2 μg of pBHRF (Schaefer et al. 2010). After transient hygromycin-resistance selection, transformants were recovered on a non-selective medium. To detect mutations, we amplified the targeting region by PCR using three sets of primers (Supplementary Table S5) and sequenced the fragments. To generate a Citrine knock-in line, the genomic sequence corresponding to intron 31-exon 35 and the 3’ flanking region of Pp*BLD10* were cloned into pCTRN-NPTII-2 plasmid (AB697058).

The Citrine-fusion plasmid was amplified by PCR using the primers listed in Supplementary Table S5 and transformed into protoplasts. Transformation for the generation of both loss-of-function mutants and Citrine knock-in lines were performed as previously described (Nishiyama et al. 2000). The Citrine knock-in lines were screened using DNA gel blot analysis to confirm single integrations. Genomic DNA (2 μg) was digested with EcoT22I, electrophoresed on 0.8% (w/v) SeaKem GTG agarose (Lonza), and transferred onto a Hybond N+ nylon membrane (GE Healthcare). Probe labelling, hybridization, and detection were performed using an AlkPhos direct labelling and detection system with CDP-Star (GE Healthcare). A PCR-amplified fragment of the 3’ untranslated region of Pp*BLD10* was used as a DNA probe.

### Microscopy

To observe Marchantia spermatids, antheridia were fixed for 60 min with 4% (w/v) paraformaldehyde (PFA) in PME buffer (50 mM PIPES-KOH, 5 mM EGTA, and 1 mM MgSO_4_ [pH 6.8]), and treated for 30 min with cell wall digestion buffer (1% [w/v] cellulase, 0.25% [w/v] pectolyase Y-23, 1% [w/v] BSA, 0.1% [w/v] NP-40, 1% glucose, and 1× cOmplete™ EDTA-free protease inhibitor cocktail [Roche Applied Science] in PME buffer). The samples were placed on a glass slide and then covered with a cover slip in PBS containing 0.1% (v/v) Hoechst33342 (Dojindo). For immunostaining of Marchantia spermatids, antheridia were fixed for 90 min with 4% (w/v) PFA in PME buffer and treated for 30 min with cell wall digestion buffer. Cells were then treated with permeabilization buffer (0.01% [v/v] Triton X-100 and 1% [w/v] BSA in PME buffer) for 10 min. After washing with PME buffer three times, cells were placed on a MAS-coated glass slide (Matsunami) and incubated for 30 min at room temperature with blocking solution (1% [w/v] BSA in PBS buffer). After removal of blocking solution, the cells were incubated with primary antibody in PBS buffer at 4°C overnight.

After washing with PBS buffer three times, the samples were incubated for 60 min at 37°C with the secondary antibody and 0.1% (v/v) Hoechst33342 in PBS buffer. After washing with PBS buffer three times, the slides were mounted using ProLong Diamond Antifade reagent (Thermo Fisher Scientific). Samples were observed under a confocal microscope (LSM780, Carl Zeiss) with an oil immersion lens (×63). For immunostaining of Physcomitrella spermatids, the same method was used with modifications. The duration of cell wall digesting buffer treatment was changed to 40 min. Samples were observed under a confocal microscope (LSM880, Carl Zeiss) with an oil immersion lens (×63).

For observation of Marchantia spermatozoids, freshly prepared spermatozoids in distilled water were observed under a dark-field microscope (Olympus) equipped with an ORCA-Flash4.0 V2 camera (Hamamatsu Photonics). Physcomitrella spermatozoids were extracted by pressing the antheridia between a glass slide and a cover slip with a 0.03-mm spacer (Koshimizu et al. 2018) and observed under a dark-field microscope (BS-2040T, BioTools) equipped with a Michrome 5Pro camera (BioTools).

The obtained images were processed using ImageJ (National Institutes of Health) and Photoshop (Adobe Systems) software.

### Antibodies

The monoclonal antibody against acetylated tubulin was purchased from Sigma-Aldrich (T7451) and used at 1/10000 dilution for immunostaining. The polyclonal antibody against centrin was described previously (Higo et al. 2018) and used at 1/5000 dilution for immunostaining. Alexa Fluor 488 plus goat anti-mouse IgG, Alexa Fluor 594 plus goat anti-rabbit IgG, and Alexa Fluor 680 goat anti-mouse IgG were purchased from Thermo Fisher Scientific and used at 1/1000 dilution for immunostaining.

### Transmission electron microscopy

To observe spermatids, antheridia of Marchantia Tak-1, the Mp*bld10-1* mutant, Physcomitrella wild type, and Pp*bld10-30* were collected and fixed with 2% PFA and 2% glutaraldehyde (GA) in 0.05 M cacodylate buffer (pH 7.4) at 4°C overnight. The fixed samples were washed three times with 0.05 M cacodylate buffer for 30 min each and then post-fixed with 2% osmium tetroxide in 0.05 M cacodylate buffer at 4°C for 3 h. The samples were dehydrated in graded ethanol solutions (50% and 70% ethanol for 30 min each at 4°C, 90% for 30 min at room temperature, four times with 100% ethanol for 30 min at room temperature, and 100% ethanol overnight at room temperature). The samples were infiltrated with propylene oxide (PO) twice for 30 min each and placed into a 50:50 mixture of PO and resin (Quetol-651; Nisshin EM Co.) for 3 h. The samples were transferred to 100% resin and polymerized at 60°C for 48 h. To observe mature Marchantia spermatozoids, spermatozoids of Tak-1 and the Mp*bld10-1* mutant were collected in water, centrifuged at 5000×*g* for 3 min, and fixed with 2% PFA and 2% GA in 0.05M cacodylate buffer (pH 7.4) at 4°C overnight. The fixed samples were washed three times with 0.05 M cacodylate buffer for 30 min each and post-fixed with 2% osmium tetroxide in 0.05 M cacodylate buffer at 4°C for 2 h. The samples were dehydrated in graded ethanol solutions (50% and 70% ethanol for 20 min each at 4°C, 90% for 20 min at room temperature, and four times with 100% ethanol for 20 min at room temperature). The samples were infiltrated with PO twice for 30 min each and placed into a 70:30 mixture of PO and resin (Quetol-651; Nisshin EM Co.) for 1 h, then the tube cap was opened, and PO was allowed to volatilize overnight. The samples were transferred to 100% resin and polymerized at 60°C for 48 h. The polymerized resins were ultra-thin-sectioned at 70 nm with a diamond knife using an ultramicrotome (Ultracut UCT; Leica) and the sections were mounted on copper grids. The sections were stained with 2% uranyl acetate for 15 min at room temperature, washed with distilled water, and secondary-stained with lead stain solution (Sigma-Aldrich) for 3 min at room temperature. The grids were observed under a transmission electron microscope (JEM-1400Plus; JEOL Ltd.) at an acceleration voltage of 100 kV. Digital images (3296×2472 pixels) were acquired using a CCD camera (EM-14830RUBY2; JEOL Ltd.).

## Acknowledgements

We thank N. Kawakami (Meiji Univ.), K. Yoshimoto (Meiji Univ.), and H. Kaku (Meiji Univ.) for support in environment of experiments, F. Nogué (INRA), Y. Horiuchi (NIBB), and M. Hasebe (NIBB) for providing plasmids for genome editing of Physcomitrella. We also thank T. Kimura (Hokkaido Univ.) and M. Shimamura (Hiroshima Univ.) for anti-centrin antibody and H. Mano (NIBB) for lending electron microscopy instrumentation and for general advice. Computations were partially performed on the NIG supercomputer at the ROIS National Institute of Genetics and the Data Integration and Analysis Facility at the National Institute for Basic Biology. Electron microscopy analyses were partly supported by the EM facility of the National Institute for Physiological Sciences. Rooms for cultivating Marchantia were provided by the Model Plant Research Facility, NIBB Bioresource Center. This work was supported by JSPS KAKENHI grants to K.Y. (19H04870), K.E. (19H04872), T.U. (19H05675 and 21H02515), N.M. (20K15824), T.N (15H04413 and 19K22448) and K.S. (18K06367).

## Author contributions

S.K. and K.Y. performed the computational analyses. N.M., K.E., and T.U. conducted the functional analyses in Marchantia. S.K., E.Y., and K.S. conducted the functional analyses in Physcomitrella. S.K. and T.N. performed sequence and phylogenetic analyses. All authors analyzed the data and participated in writing the manuscript.

