## Supplementary Figures for "Phylogenetic distribution and expression pattern analyses identified a divergent basal body assembly protein involved in land plant spermatogenesis"

**Step1, Selection the protein family groups specific in the plant species of producing spermatozoid**

**935 protein family groups**

| | Species | Group 1 | Group 2 | Group 3 | … |
|---|---|---|---|---|---|
| Plants producing spermatozoid | *Ginkgo biloba* | ○ | ○ | - | |
| | *Selaginella moellendorffii* | ○ | ○ | - | |
| | *Marchantia polymorpha* | ○ | ○ | - | |
| | *Physcomitrium patens* | ○ | ○ | - | |
| | *Chara braunii* | ○ | ○ | - | |
| | *Chlamydomonas reinhardtii* | ○ | - | - | |
| Animals | Human | ○ | - | ○ | |
| | Mouse | ○ | - | ○ | |

**Step2, Extraction of the protein family groups composed by the genes highly expressed during spermatogenesis**

**165**

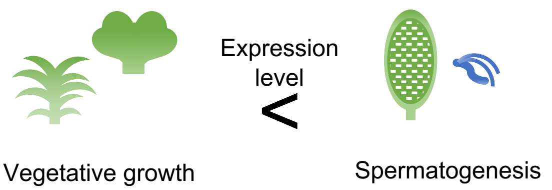


**Step3, Selection of the protein family groups consisting of the members showing low BLAST similarities with animals and Chlamydomonas**

**31**

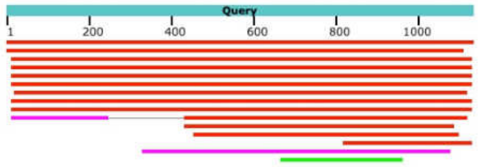


**Step4, Human check of the expression level data in Step2**

**4**

- Lower expression levels in vegetative growth stage
- Substantial difference in expression levels between vegetative and male reproductive stages

**Step5, Final selection by known functional information**

**2**

Group-89 and Group-230

Supplementary Fig. S1. A workflow of *in silico* analysis. Orange circles show the number of remain protein family groups by selections.

A

| | |
|---|---|
| WT | TGAAATTTCCCATGACCTACGAGTCAAAGTTGCAGCCCTCA**CGG**ACAGTCTGAAGCAG |
| Mp*bld10-2* | TGAAATTTCCCATGACCTACGAGTCAAAGTTGCAGCCT------------------- |
| WT | GAGAGACTGCGCGTGGAACAACTTGTCTGCGAGAATCAGAAGCTTGTTAAAGATAAAG |
| Mp*bld10-2* | ---------------------------------------------------------- |
| WT | CGGAACTCGAAAAAAATGCCCAAGAATTCAGAGATTTGATCAGCGAATTAAAAAAACA |
| Mp*bld10-2* | ---------------------------------------------------------- |
| WT | TGAAAACGAAGAAGAGCAAGTGCAACAGTGGAGAGCACAAAGTATGCATCAGCTCCAG |
| Mp*bld10-2* | ---------------------------------------------------------- |
| WT | gcaagatatcttcccttcatcctataacacgctactgcaggccatagagtcgttcaat |
| Mp*bld10-2* | ---------------------------------------------------------- |
| WT | ccagcttccgagtcgtttctccatcgacatcctcgtttagtggggcgtggaactggta |
| Mp*bld10-2* | ---------------------------------------------------------- |
| WT | tccagtacgaaaccatcgactccaggtccgtaaatgacaactagtccacggaaggatc |
| Mp*bld10-2* | ---------------------------------------------------------- |
| WT | tgccgctgtgcagctagtaaatgtatcacttgaaatcttaggaagatgacccaataac |
| Mp*bld10-2* | ---------------------------------------------------------- |
| WT | tgggagtgcacctttaccttcgtgatcatgacaccacgt |
| Mp*bld10-2* | -----gtgcacctttaccttcgtgatcatgacaccacgt |

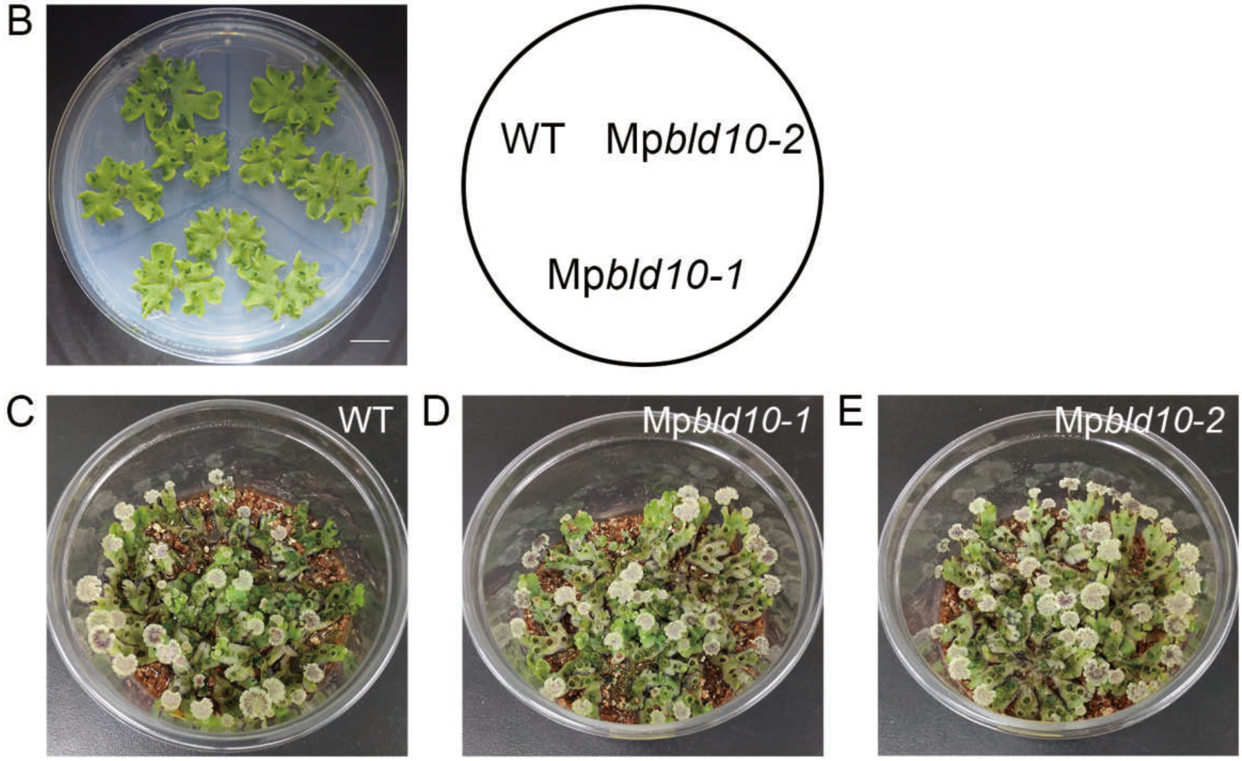


Supplementary Fig. S2. Vegetative growth and antheridiophore formation were not affected by Mp*bld10-1* and Mp*bld10-2* mutations. (A) Nucleotide and amino acid sequences around mutated sites in wild-type and Mp*bld10-2* are aligned. The target and PAM sequences are indicated underlined and in bold, respectively. (B) An image of 15-day-old gemmalings of wild-type, Mp*bld10-1*, and Mp*bld10-2* cultured on an agar plate of 1/2 Gamborg B5 medium. (C-E) Images of thalli of wild-type (B), Mp*bld10-1* (C), and Mp*bld10-2* (D) with antheridiophores. Scale bars = 1cm.

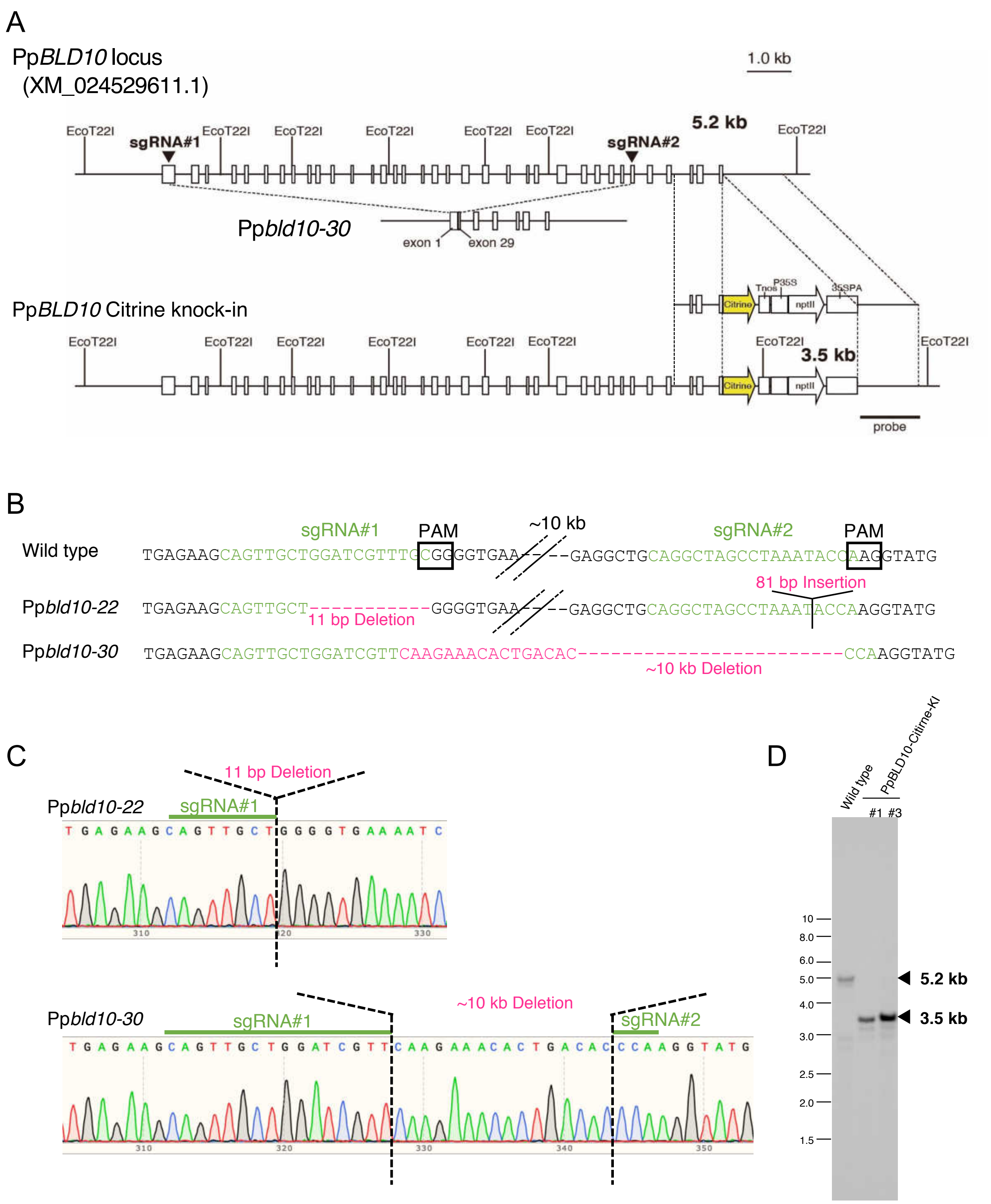


Supplementary Fig. S3. Generation of CRISPR-Cas9-mediated mutants and Citrine knock-in lines of Pp*BLD10*. (A) A schematic representation of Pp*BLD10* locus on Physcomitrium genome. The white boxes indicate exons and the black triangles indicate sgRNA-targeted sequences. Yellow arrow, white arrow, and boxes indicate indicate yellow fluorescent protein gene *Citrine*, the neomycin phosphotransferase II gene (nptII), the terminator of nopaline synthase gene (Tnos), the modified promoter of cauliflower mosaic virus 35S (P35S), the polyadenylation signal of cauliflower mosaic virus 35S (35SPA) on the pCTRNNPTII-2 plasmid (AB697058), respectively. (B, C) The results of sequences of sgRNA-targeted sites in wild type and the CRISPR mutants (Pp*bld10-22* and Pp*bld10-30*). The sgRNA-targeting sequences, mutated sequences, and PAM indicate in green letters, magenta letters, and black boxes, respectively. (D) DNA gel-blot analysis of Citrine knock-in lines. Genomic DNAs were digested with *Eco*T22I.

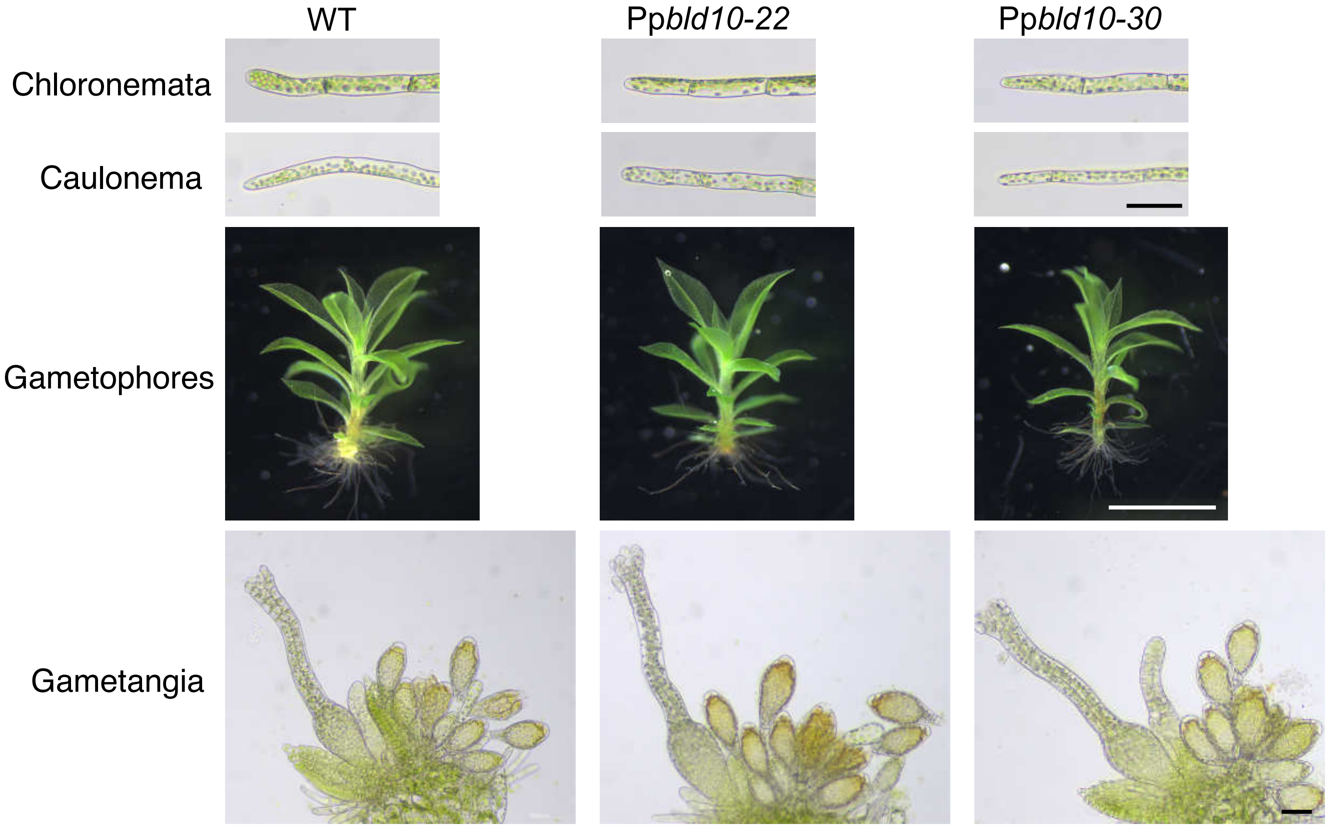


Supplementary Fig. S4. No difference was detect between wild type and Pp*bld10-22* or Pp*bld10-30* mutants. Representative images of the chloronema filaments, caulonema filaments, gametophores, and antheridia and archegonia in the wild type, Pp*bld10-22*, and Pp*bld10-30* mutants. Scale bars = 50 μm in chloronema filaments, caulonema filaments, and antheridia and archegonia; 2 mm in gametophores.

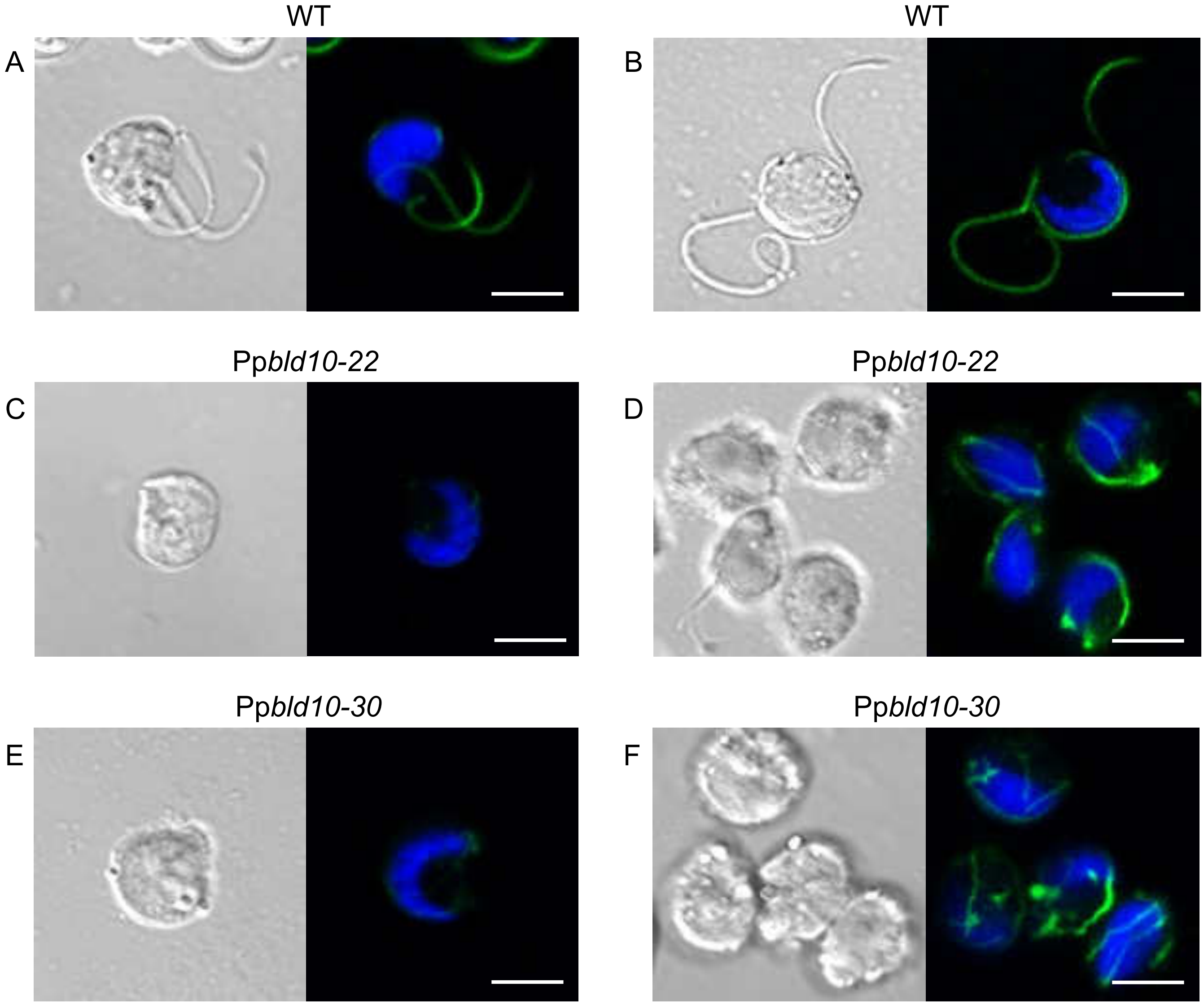


Supplementary Fig. S5. Pp*bld10* mutants have defects in flagellar formation. Maximum intensity projection images of spermatids of wild type (A and B), Pp*bld10-22* (C and D), and Pp*bld10-30* (E and F) immunostained with anti-acetylated tubulin antibodies. Nuclei were also visualized by Hoechst. Blue and green pseudo colors indicate Hoechst and acetylated tubulin, respectively. Scale bars = 5 μm.

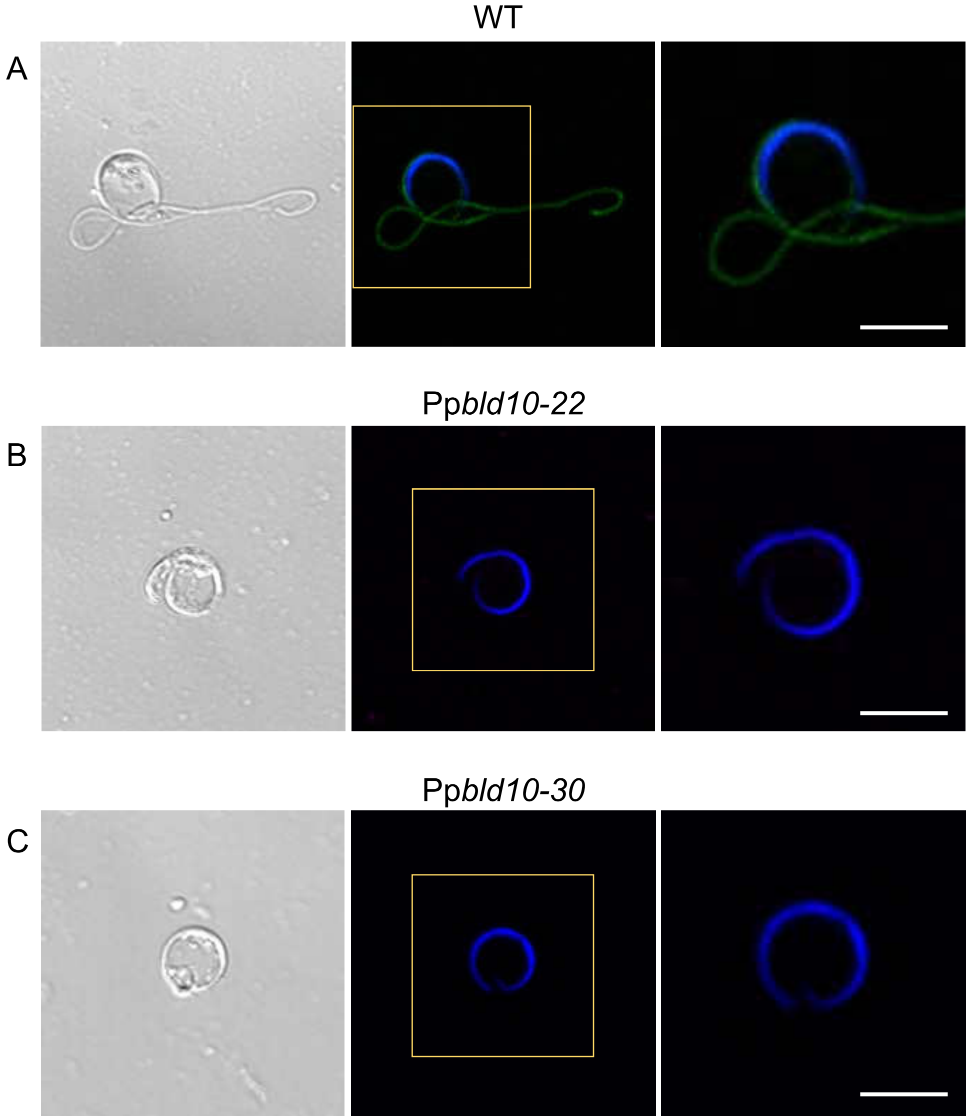


Supplementary Fig. S6. Mature spermatozoid in wild type and Pp*bld10* mutants. Maximum intensity projection images of spermatozoid of wild type (A), Pp*bld10-22* (B), and Pp*bld10-30* (C) immunostained with anti-acetylated tubulin antibodies. Nuclei were also visualized by Hoechst. Blue and green pseudo colors indicate Hoechst and acetylated tubulin, respectively. Scale bars = 5 μm.

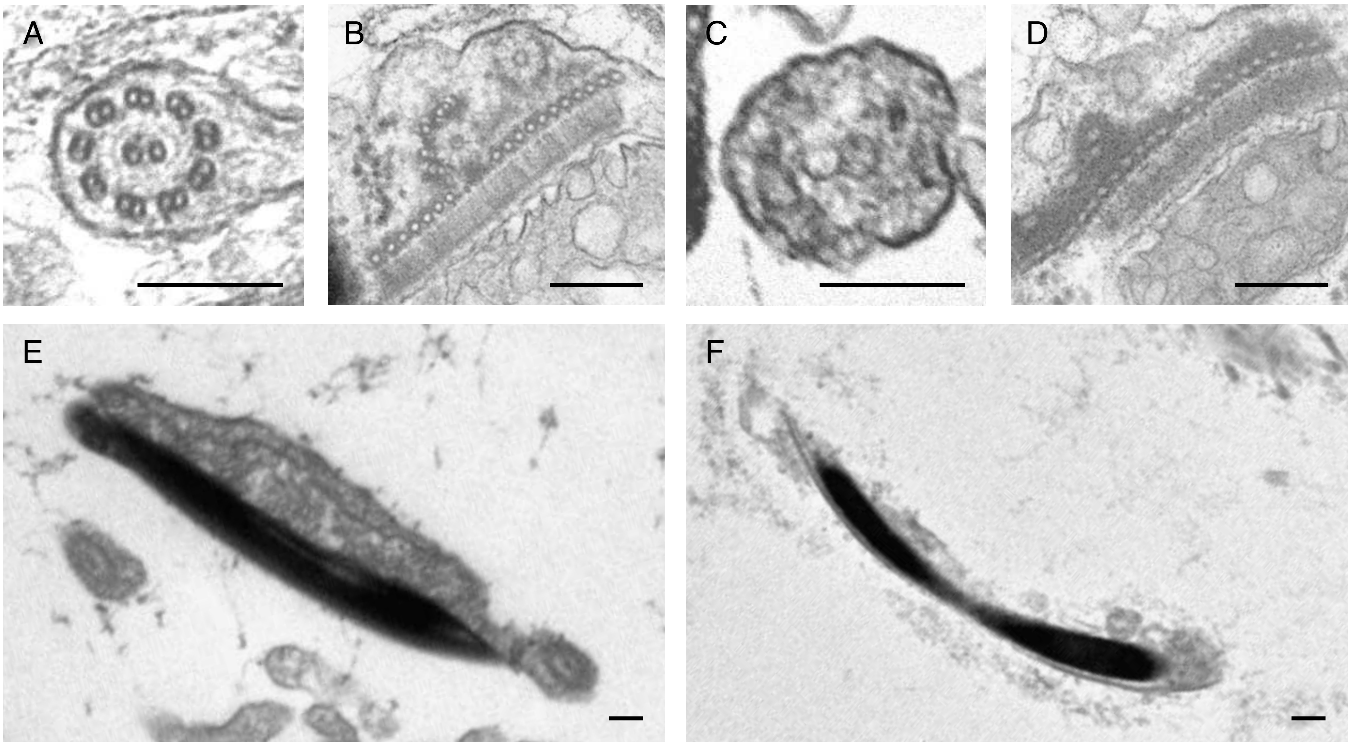


Supplementary Fig. S7. Transmission electron microscopy (TEM) of spermatids and spermatozoids in wild type and Pp*bld10-30*. (A-D) TEM images in spermatids of wild type (A and B) and Pp*bld10-30* (C and D). Axonemes in flagella (A and C), multilayered structures and basal bodies (B and D) are shown. (E and F) TEM images of nuclei in spermatozoids of wild type (E) and Pp*bld10-30* (F). Scale bars = 200 nm.

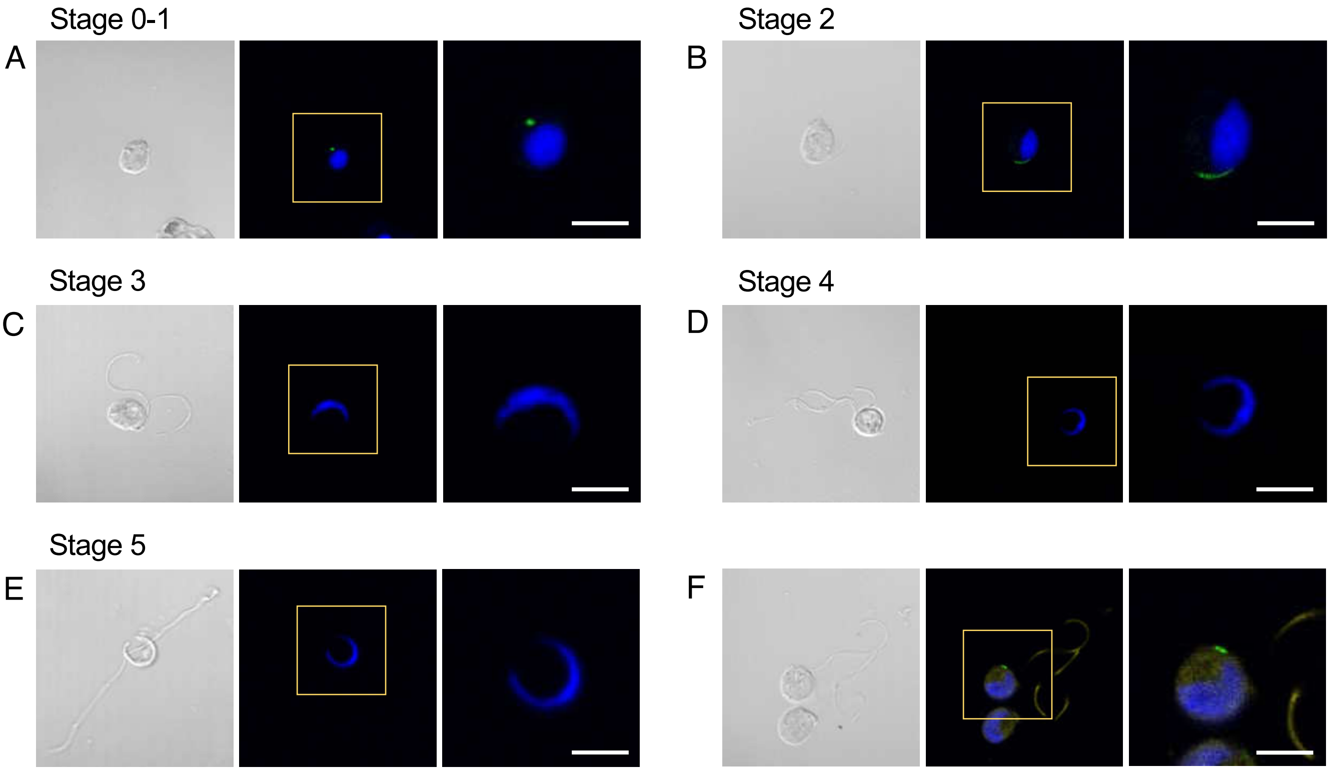


Supplementary Fig. S8. Subcellular localization of the PpBLD10 protein. (A-E) DIC and maximum intensity projection images of spermatids and spermatozoids expressing PpBLD10-Citrine (green) at stage 0-1 (A), stage 2 (B), stage 3 (C), stage 4 (D), and stage 5 (E), whose nuclei were also stained with Hoechst (blue). Developmental stages are classified according to Minamino *et al*. (2021). (F) Maximum intensity projection images of a spermatid expressing PpBLD10-Citrine (green) immunostained with anti-acetylated tubulin (yellow) antibodies. The nucleus was stained with Hoechst (blue). Scale bars = 5 µm.

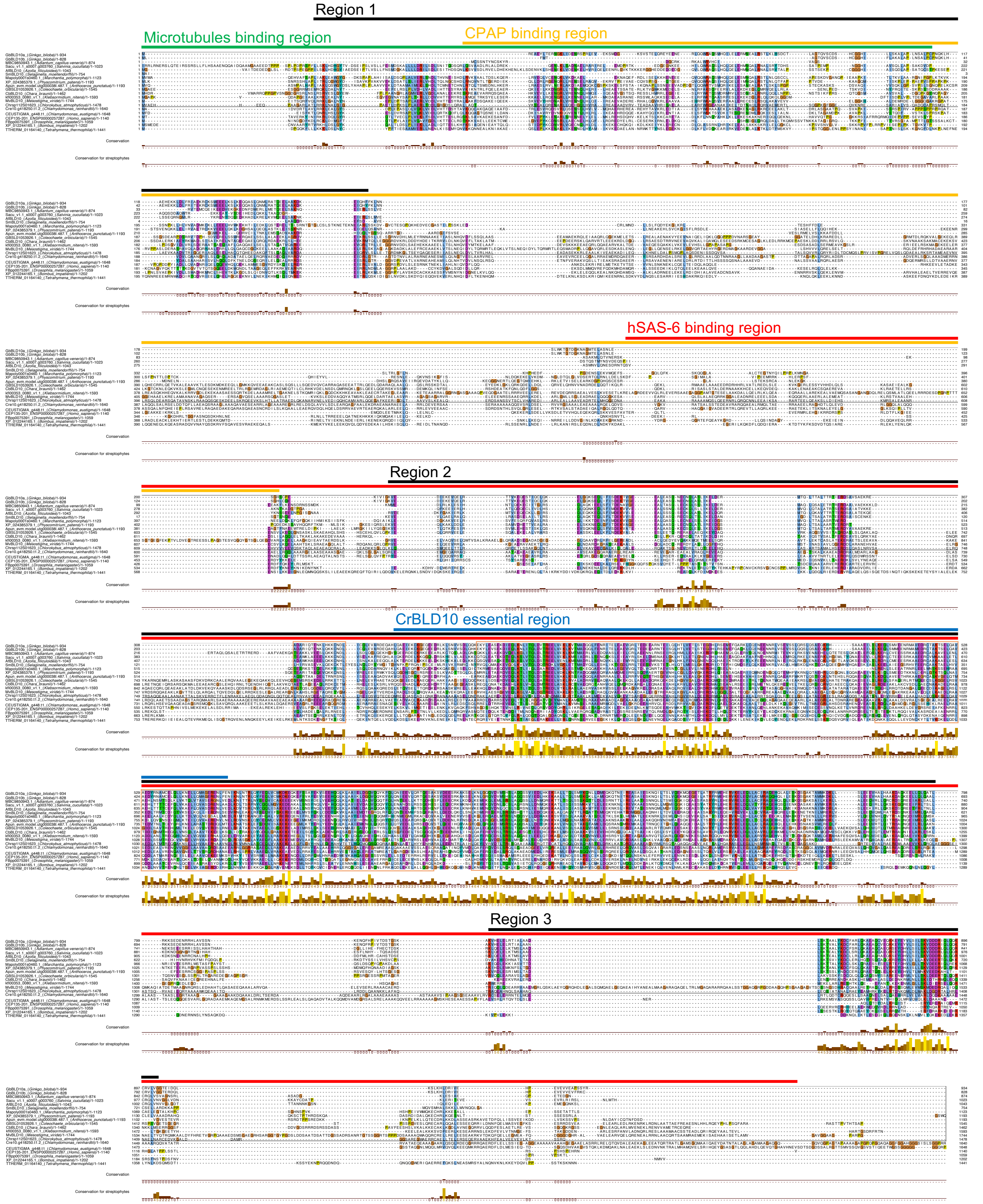


Supplementary Fig. S9. Multiple alignment of BLD10/CEP135 family proteins. From the top, sequences of Ginkgo (a and b), Adiantum, Salvinia, Azolla, Selaginella, Marchantia, Physcomitrium, Anthoceros, Coleochaete, Chara, Klebsormidium, Mesostigma, Chlorokybus, *C. reinhardtii*, *C. eustigma*, human, fly, bee, and Tetrahymena. Histograms show conservation levels among all those sequences (top) and streptophytes (bottom). Green, yellow, red, and blue lines indicate the microtubules binding region, CPAP binding region, hSAS-6 binding region, and the probably essential region of CrBLD10, respectively. Black lines indicate each conserved region in the streptophytes. Orange boxes indicate regions using construction of phylogenetic tree and molecular clock analyses.
